# Weighted Centroid Trees: A general approach for summarizing phylogenies in tumor mutation tree inference

**DOI:** 10.1101/2023.09.11.557167

**Authors:** Hamed Vasei, Mohammad Hadi Foroughmand Araabi, Amir Daneshgar

## Abstract

Tumor mutation trees are the primary tools to model the evolution of cancer. Not only some tumor phylogeny inference methods may produce a set of trees having potential and parallel evolutionary histories, but also mutation trees from different patients may also exhibit similar evolutionary processes. When a set of correlated mutation trees is available, compressing the data into a single best-fit tree, exhibiting the shared evolutionary processes, is definitely of great importance and can be beneficial in many applications. In this study, we present a general setup to study and analyse the problem of finding a best-fit (centroid) tree to a given set of trees and we use our general setup to analyse mutation trees as our main motivation. For this let *ε* : *𝒯*_*n*_ → ℝ^*n*×*n*^ be an embedding of labeled rooted trees into the space of real square matrices and also let *L* be a norm on this space. We introduce the *nearest mapped tree* problem as the problem of finding a closest tree to a given matrix **A** with respect to *ε* and *L*, i.e., a tree *T* ^*^(**A**) for which *L*(*ε*(*T* ^*^(**A**)) − **A**) is minimized. Within this setup, our potential candidates for the embedding are *adjacency, ancestry*, and *distance* matrices of trees, where we consider the cases of *L*_1_ and *L*_2_ norms in our analysis. We show that the function d(*T*_1_, *T*_2_) = *L*(*ε*(*T*_1_) − *ε*(*T*_2_)) defines a family of dissimilarity measures, covering previously studied *parent-child* and *ancestor-descendent* metrics. Also, we show that the nearest mapped tree problem is polynomial-time solvable for the adjacency matrix embedding and is *𝒩𝒫*-hard for the ancestry and the distance embeddings. The *weighted centroid tree problem* for a given set of trees of size *k* is naturally defined as a nearest mapped tree solution to a weighted sum of the corresponding matrix set. In this article we consider uniform weighted-sums for which all weights are equal, where in particular, the (classical) *centroid tree* is defined to be a solution when all weights are chosen to be equal to 1*/k* (i.e., the mean case). Similarly, the *ω*-weighted centroid tree is a solution when all weights are equal to *ω/k*. To show the generality of our setup, we prove that the solution-set of the centroid tree problem for the adjacency and the ancestry matrices are identical to the solution-set of the *consensus tree problem* for parent-child and ancestor-descendent distances already handled by the algorithms GraPhyC(2018) and TuELiP(2023), respectively. Next, to tackle this problem for some new cases, we provide integer linear programs to handle the nearest mapped tree problem for the ancestry and the distance embeddings, giving rise to solutions of the weighted centroid tree problem in these cases. To show the effectiveness of this approach, we provide an algorithm, <u>WAncILP_2_</u>, to solvethe 2-weighted centroid tree problem for the case of the ancestry matrix and we justify the importance of the weighted setup by showing the pioneering performance of <u>WAncILP_2_</u> both in a comprehensive simulation analysis as well as on a real breast cancer dataset, in which, by finding the centroids as representatives of data clusters, we provide supporting evidence for the fact that some common aspects of these centroids can be considered as suitable candidates for reliable evolutionary information in relation to the original data. metrics.

## 1 Introduction

Cancer is often modeled as an evolutionary stochastic process that explains inter- and intra-tumor heterogeneity [FV90; MP10]. Despite the heterogeneity, there are hallmarks of cancer, characterized by repeated evolutionary events and behaviors [HW11]. Heterogeneity necessitates the study of individual patients using their own data, however, high-throughput data sequencing is often noisy and may contain many random artifacts [HBD10; Par+16]. On the other hand, the recurring patterns in cancer evolution allow one to utilize data from different patients, reduce the noise of each single data-source, and infer the shared evolutionary histories more accurately.

REVOLVER [Car+18] is an influential method incorporating multi-patient data, which uses transfer learning to jointly infer trees for a set of patients. REVOLVER iteratively compares inferred trees as partial orderings of the mutations and modifies them through an expectation maximization procedure. HINTRA [Kha+19] uses the same idea as REVOLVER but considers more distant relationships among mutations in a tree at the cost of increasing the run-time complexity. CONETT [Hod+20] uses ancestral relationships between mutations in different trees to infer a single tree that depicts conserved evolutionary trajectories, by solving a combinatorial optimization problem. Another method to compute significant evolutionary trajectories from a set of phylogenetic trees is MASTRO [PV22], which assesses the significance of the trajectories using a conditional statistical test that captures the coherence in the order in which alterations are observed in different tumors. TreeMHN [LKB23], provides probabilistic measures of potential evolutionary trajectories and patterns of clonal exclusivity or co-occurrence from a cohort of intra-tumor phylogenetic trees.

By summarizing a set of trees, one intends to find a tree that, “in the mean”, may be considered as a legitimate representative of the set. Naturally, formulating the term “in the mean” gives rise to applying the theory of metrics and distance measures to the subject. Usually, distance measures in classical phylogeny are defined for leaf-labeled phylogenies. A popular and classical distance is the Robinson-Foulds metric [RF81] which compares bi-partitions induced by two phylogenetic trees and sums up the number of their unique bi-partitions. Also, recently, several dissimilarity measures on fully labeled trees have been proposed. The MLTD [Kar+19], permutation, and rearrangement distances [Ber+19; BBG20] are edit distances between mutation trees. Bourque distance [JBZ21], the similarity measure used in ConTreeDP algorithm [FS21], and the generalized Robinson-Foulds distance for phylogenetic trees [LRV20; LRV21] are also defined through comparing bi-partitions. MP3 [Cic+21] is a similarity measure that compares mutation triplets of the two trees. L. Oesper *et al*. introduced parent-child distance (PCD), ancestor-descendant distance (ADD), common ancestor set distance (CASet), and distinctly inherited set comparison distance (DISC) metrics by looking at the ancestry relations in mutation trees [GSO18; DiN+20].

Consensus trees are also well-known objects in phylogenetics to summarize a set of trees [Bry03]. GraPhyC [Gov+20] is the first method to compute a consensus tree for a set of mutation trees by minimizing the total parent-child distance. ConTreeDP [FS21] uses bi-partitions to find a consensus tree, while TuELiP [GSO23] finds a consensus tree based on ancestral relationships among mutations (ancestor-descendent distance). Note that finding a consensus tree with respect to the ancestor-descendant distance is an -hard [QE23] problem. The *Multiple Consensus Tree* [AQE19] and the *Multiple Choice Consensus Tree* [Chr+20] problems, generalize the consensus tree problem to simultaneously cluster trees and infer a consensus tree for each cluster.

PhyC, [Mat+17] is an algorithm that uses embedding and metrics to cluster evolutionary trees. This algorithm first creates a large reference tree by combining data from a set of patients. Then, by defining a mapping between individual trees of patients and the reference tree, a numerical vector is assigned to each individual tree. These vectors are used in an Euclidean setting to apply hierarchical clustering.

The goal of this study is to provide a general framework that may be used to summarize/compress a set of trees into a single tree. Our approach consists of three major steps: i) mapping the given trees into the space of real matrices, ii) finding the centroid point of these mapped images (as a mean/average matrix), and then iii) finding the best tree-approximation of the centroid matrix as a labeled tree. For this, one naturally needs an *embedding* map for the first step and a *distance* for the averaging process. The last step is actually an optimization problem asking for the closest point in the space of labeled trees to the average matrix obtained through the averaging process, which, hereafter, is referred to as the *nearest mapped tree* problem.

In what follows, we analyse this general setup for a variety of cases including adjacency, ancestry, and distance matrix-embedding of trees and entry-wise *L*_*p*_ norms (when *p υ* {1, 2}). Within this setup, we define a family of dissimilarity measures between mutation trees as the norm of the difference between their mapped images, where this reveals that previously studied *consensus tree problems* solved by GraPhyC and TuELiP fit into our generalized framework.

Our general framework not only makes it possible to use a variety of embeddings and norms, but also it is flexible in applying *weighted sum* operators to obtain centroids. In particular, we introduce the algorithm WAncILP_2_ and we show that it improves the weighting strategy used in TuELiP. We also provide a generalized simulation setup containing different possible tree alternations and evaluate our algorithms through a comprehensive analysis in comparison to other existing algorithms, both on simulated and real data. For clarity and brevity, all the proofs and supplementary material are provided separately in the “Supplementary Materials“.

## 2 Material and methods

This section contains the definition of our general setup and the statements of our main theoretical results. In what follows, after fixing our notations we introduce the *weighted centroid tree problem* and analyse its basic properties with respect to adjacency, ancestry and distance embeddings. After that we show that the whole problem may be reduced to solving the *nearest mapped tree problem*, where we prove hardness results and provide algorithms when applicable.

### 2.1 Preliminaries and notations

Capital boldface letters (e.g. **A**) represent matrices. The element at the *i*th row and the *j*th column of a matrix **A** is denoted by **A**[*i, j*]. Given a graph *G*, the symbols *V* (*G*), *E*(*G*), and **adj**(*G*) stand for the vertex set, the edge list, and the adjacency matrix of *G*, respectively. In the sequel, all *n*-vertex graphs are vertex-labeled by the set ℤ_*n*_ = {0, …, *n-*1} where we always assume that the rows and columns of any matrix assigned to *G* are ordered according to the natural order on ℤ_*n*_ (see Fig. 1).

**Figure 1.**
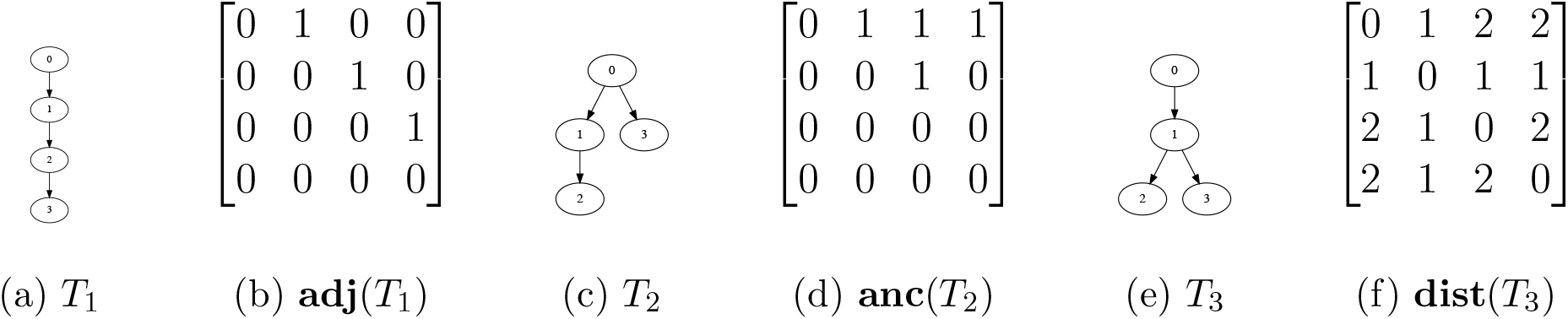
A couple of rooted trees and their matrices. (a) and (b) Mutation tree *T*_1_ and its adjacency matrix. (c) and (d) Mutation tree *T*_2_ and its ancestry matrix. (b) and (f) Mutation tree *T*_3_ and its distance matrix.

The distance between two vertices of a directed graph *G*, is the length of a shortest path between these vertices in an undirected sense. The *distance matrix* of *G*, denoted by **dist**(*G*), is defined as the matrix whose [*i, j*]th entry is the distance between the vertices *i* and *j*.

Given a directed graph *G*, a vertex *v* is the *ancestor* of a vertex *u* (or *u* is the *descendent* of *v*) if there exists a directed path from *v* to *u* in *G*. The *ancestry matrix* of *G*, denoted by **anc**(*G*), is defined as the binary matrix whose [*i, j*]th entry is 1 if the vertex *i* is the ancestor of the vertex *j*.

A *rooted tree* is a directed tree with a specific *root* vertex which is the ancestor of any other vertex. Note that directions in a rooted tree are by definition forced by the choice of the root. Because the labels in this context represent mutations, a labeled rooted tree is referred to as a *mutation tree*. The set of all *n*-vertex mutation trees is denoted by *𝒯*_*n*_.

In what follows, *ε* : *𝒯*_*n*_ → ℝ^*n×n*^ is an embedding of mutation trees into the space of real square matrices and also *L* : ℝ^*n×n*^ → R stands for a norm on this space. Fixing a weight vector **w** ∈ ℝ^*k*^, the weighted-sum operator *μ*_*ε*_ applied to a set of trees as 𝒮 = {*T*_1_, …, *T*_*k*_} is defined as

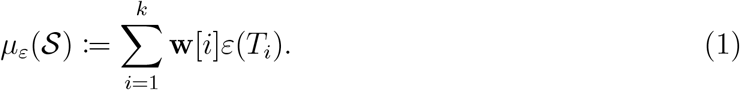

Hereafter, we assume that the weight vector is always clear from the context. Although our algorithms are general and may be applied for any weight vector, in our applications the weight vectors are always uniform with equal weights.

Also, let us recall the definition of the entry-wise *L*_*p*_ matrix norms as

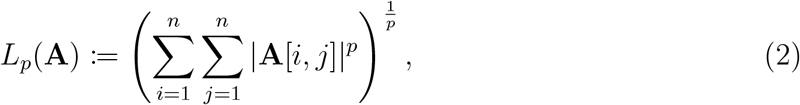

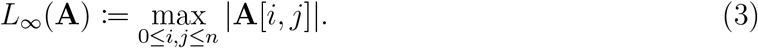

### 2.2 Dissimilarity measures

Applying our embedding setup, it is clear that

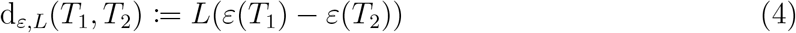

defines a dissimilarity measures on the space of trees through embeddings. Also, note that the injectivity of *ε* is a necessary and sufficient condition for d_*ε*,*L*_ to be a metric (see Supplementary Proposition 1).

**Example 1**. For the trees depicted in Fig. 1 we have

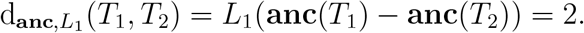

It can be verified that all dissimilarity measures defined by these embeddings and norms are computable in *𝒪* (*n*^2^) time (Supplementary Proposition 2). Also, it is interesting to note that for the parent-child distance (PCD) and the *ancestor-descendent distance* (ADD) [GSO18] we have

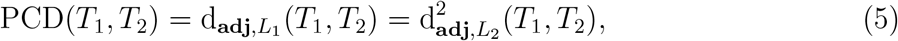

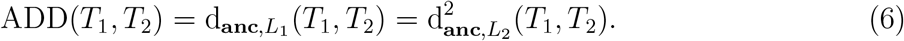

#### 2.2.1 Metrics based on the distance matrix

As opposed to the adjacency matrix, which considers local relationships, the distance matrix incorporates long-distance relationships between vertices as well as local distances. Ancestry relations are not local, but their binary nature limits their power in expressing relations between vertices. Dissimilarity measures based on **dist** mapping assess how much two trees agree with each other on distances between different pairs of elements. In other words, they measure whether labels that are close to (or far from) each other in one tree, are also close to (or far from) each other in the other tree.

Another perspective on d_**dist**,*L*_ is looking at bi-partitions of the graph. The distance between two vertices *v* and *u* is actually the number of bi-partitions of the tree in which these two vertices are in different partitions. Therefore, by comparing distance matrices of two trees, one is comparing a representation of their bi-partitions.

### 2.3 The centroid tree problem

Consensus trees, already being studied in the literature, are best-fit trees defines as follows.

#### Definition 1

(The Consensus Tree Problem (CoTP_*d*_)). Let *d* : *𝒯*_*n*_ *× 𝒯*_*n*_ → ℝ be a distance measure and set 𝒮 = {*T*_1_, …, *T*_*k*_} ⊂ *𝒯*_*n*_ be a set of mutation trees. Then the *consensus tree* of 𝒮, denoted by *T*_con_ ∈ *𝒯*_*n*_, is defined as

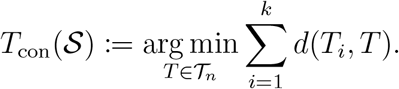

GraPhyC [Gov+20] is a polynomial time algorithm to solve CoTP_PCD_ and TuELiP [GSO23] is an ILP that solves CoTP_ADD_.

Now, following the idea of embedding trees into an Euclidean space of matrices, we introduce the a very general centroid tree as follows.

#### Definition 2

(The Weighted Centroid Tree Problem (CeTP_*ε*,*L*,*μ*_)). Let *ε* : *𝒯*_*n*_ → ℝ^*n×n*^ be an injective embedding, *L* : ℝ^*n×n*^→ ℝ be a norm, and *μ*_*ε*_ be a weighted-sum operator. Given a set 𝒮 = {*T*_1_, …, *T*_*k*_} ⊂ *𝒯*_*n*_ of mutation trees, the *weighted centroid tree* of 𝒮, denoted by *T* ^*^ ∈ *𝒯*_*n*_, is defined as

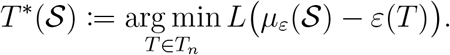

The *ω-weighted centroid tree* problem denoted by *ω*-CeTP_*ε*,*L*_ is the special case where *μ*_*ε*_ is the uniform weighted-sum with all weights equal to 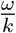. Hereafter, the problem 1-CeTP_*ε*,*L*_ is the (classical) *centroid tree* problem and is denoted by CeTP_*ε*,*L*_ in short.

The following example shows that CeTP and CoTP may have different solutions.

**Example 2**. Consider the trees *T*_1_, *T*_2_, and *T*_3_ depicted in Fig. 1. Both *T*_1_ and *T*_2_ are optimal solutions of 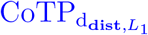 for the set 𝒮 = {*T*_1_, *T*_2_}, having the cost *L*_1_(**dist**(*T*_1_) – **dist**(*T*_2_)) = 8. To see that *T*_3_ is not an optimal consensus tree note that,

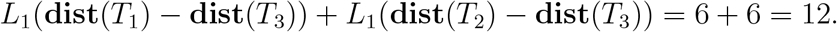

On the other hand, *T*_3_ is the single optimal solution of 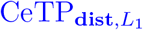 having the cost

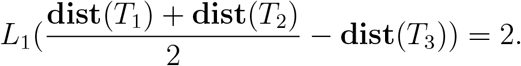

None of *T*_1_ and *T*_2_ is an optimal centroid tree since for any *T* ∈ 𝒮,

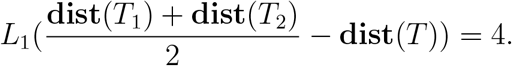

The next theorem demonstrates some cases when the solution sets of these optimization problems are equal.

#### Theorem 1.

Given any set of mutation trees *S* ⊂ *𝒯*_*n*_,

a. for any binary embedding *ε* : *𝒯*_*n*_ → {0, 1}^*n×n*^, the solution sets of 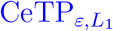 and 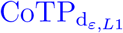 are identical.
b. for any embedding *ε* : *𝒯*_*n*_ → ℝ^*n×n*^, the solution sets of 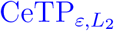 and 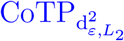 are identical.

By the fact that both **adj** and **anc** are binary embeddings, one deduces that,

#### Corollary 1.

The following sets of optimization problems admit the same set of solutions:

a. 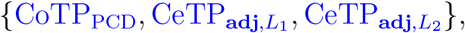,
b. 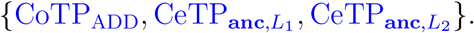.

### 2.4 The nearest mapped tree problem

Our main observation is the fact that the weighted centroid problem may be reduced to a general *nearest tree* problem as follows. In the rest of this section we will concentrate on this problem and provide some complexity results as well as some algorithms to solve it in some important cases.

#### Definition 3.

(The Nearest Mapped Tree Problem (NMTP_*ε*,*L*_)). Let *ε* : *𝒯 𝒯*_*n*_ ℝ^*n×n*^ be an embedding and *L* : ℝ^*n×n*^ → ℝ be a norm. Given a matrix **M** ∈ ℝ^*n×n*^, a nearest mapped tree *T*^*^ (**M**) is defined as

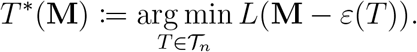

Actually, the nearest mapped tree problem for the adjacency embedding is easy to solve.

#### Theorem 2.

Given a real *n × n* matrix *M*,

a. the problem 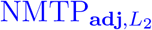 can be solved in *𝒪* (*n*) time through finding a maximum weight spanning arborescence tree.
b. the problem 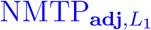 can be solved in *𝒪* (*n*^3^) time by TrimArb algorithm (Supplementary Algorithm 2).

For the ancestry matrix embedding, recently, [QE23] proved that CoTP_ADD_ is an NP-hard problem. Hence, Corollary 1 implies that NMTP_**anc**,*L*_ is also *𝒩𝒫*-hard.

#### Corollary 2.

The problems 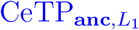, 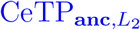, 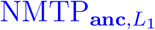, and 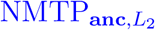 are *𝒩𝒫*-hard.

On the algorithmic side, we developed two integer linear programs (ILP) solving 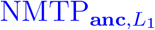 and 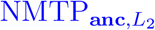, namely AncL1 (Supplementary Fig. 8) and AncL2 (Supplementary Fig. 7), respectively. These ILPs have the same set of constraints as TuELiP, but their cost functions are quite different.

For the distance matrix embedding we first prove the following hardness result.

#### Theorem 3.

The problems 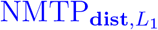 and 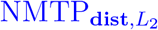 are *𝒩𝒫*-hard.

On the algorithmic side, we provide the ILP program DistL1 (see Fig. 2) to solve the problems 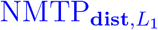 and 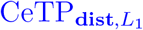. Note that the problem 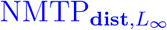 can also be solved similarly by a minor modification of DistL1, named DistLInf (Supplementary Fig. 10).

**Figure 2.**
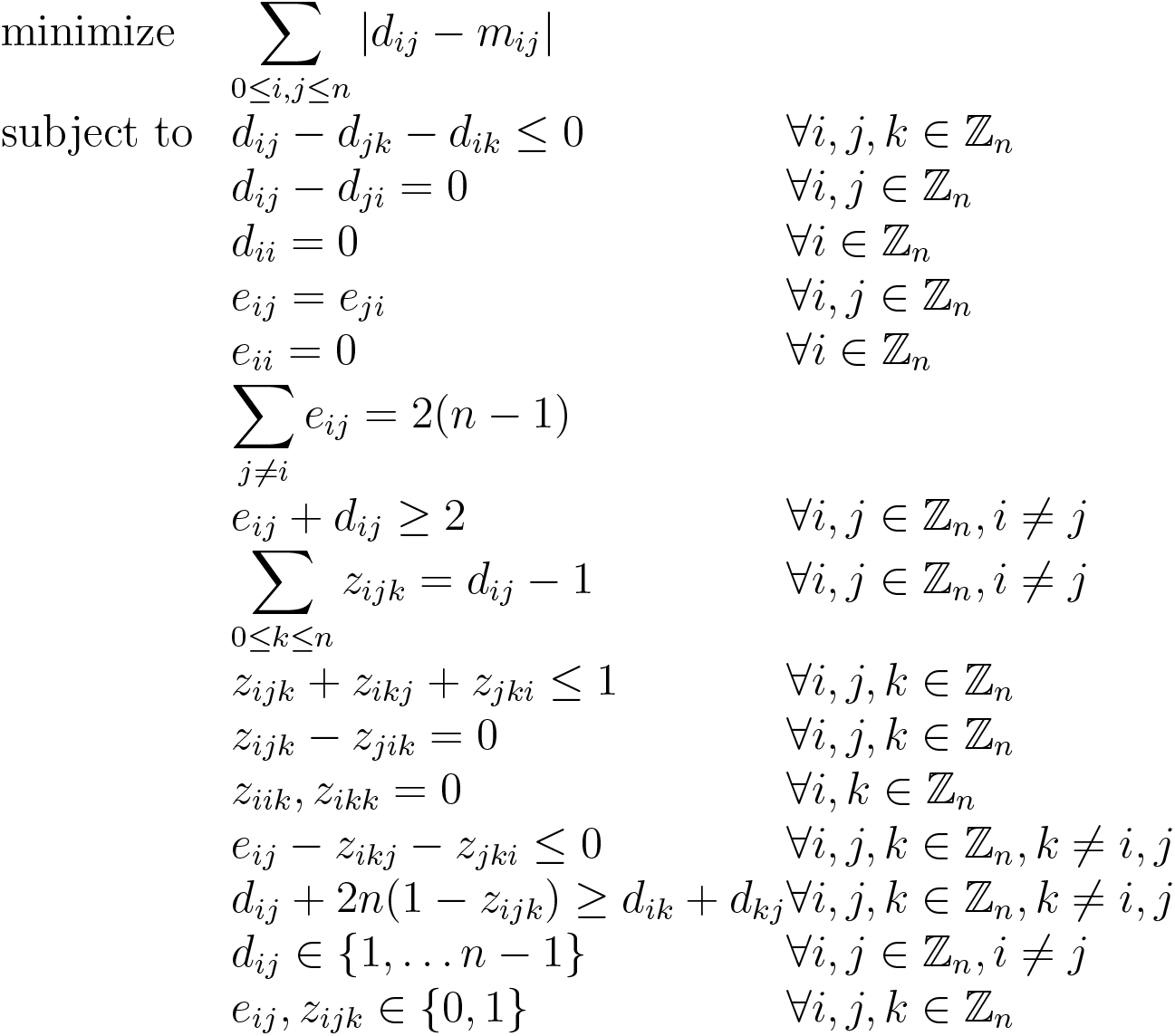
The ILP program DistL1 that solves 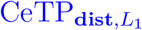. The input is the **M** having *m*_*ij*_ as its entries. For an optimal solution for this ILP, the matrix **D** having *d*_*ij*_ as its entries represents **dist**(*T*), the matrix **E** represents **adj**(*T*) and for all *i, j, k*, the condition *z*_*ijk*_ = 1 indicates that vertex *k* is on the path between *i* and *k* in *T*.

#### Theorem 4.

Given a matrix **M** ∈ ℝ^*n×n*^, the ILP program DistL1 solves 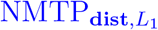.

### 2.5 A second look at weighted centroids

Note that, in general, one may apply weights to compute centroids, in order to summarize a set of tumor trees more efficiently. Regarding adjacency matrices, since the maximum weight spanning arborescence tree does not change by scaling, we have the following corollary, indicating that the case of adjacency matrix embedding is not interesting when one is concerned with weighted centroids.

#### Algorithm 1: WAncILP_ω_

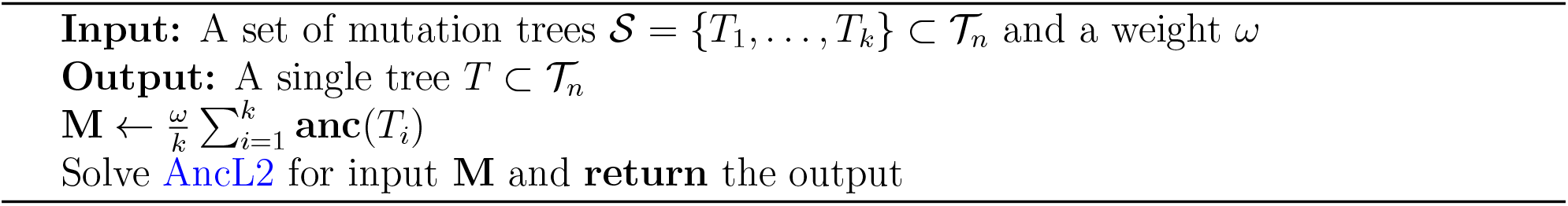

#### Corollary 3.

For any *ω >* 0 the solutions to the problems 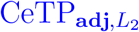 and 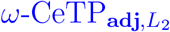 are identical.

On the contrary, using weighted setups has already been proved to be useful for the ancestry matrix embedding, in TuELiP, that applies weights on the input trees. From this point of view, our weighted centroid tree problem may be considered as a generalization of the *weighted consensus tree* problem solved by TuELiP (Supplementary Theorem 14).

Note that, in the context of ancestry relations, one could argue that the signals are asymmetric, meaning that the presence of ancestry signals is more informative than their absence. For example, if mutation *m*_1_ is observed as an ancestor of mutation *m*_2_ in 49 out of 100 patients, this is a strong indication that *m*_1_ is an ancestor of *m*_2_. However, CeTP_**anc**,*L*_ treats the presence and absence of ancestry relations equally. In this case, **M**_*c*_[*m*_1_, *m*_2_] = 0.49 for the centroid point, while the absence of an ancestry relation is preferred when solving the centroid tree problem.

Assuming that one does not have any extra evidence for choosing the weights, to increase the weight of ancestry signals one may solve 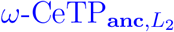 for some candidate values of *ω >* 1 using the WAncILP_*ω*_ algorithm (Algorithm 1). According to our example having 49 observation of *m*_1_ as an ancestor of *m*_2_, one may verify that for *ω* = 2 the new value for **M**_*c*_[*m*_1_, *m*_2_] is 0.98 which is quite close to 1. We have tried *ω* = 2 and *ω* = 3 in our analysis where results show that *ω* = 2 seems to be a better choice, however, the best choice for *ω* may not be an integer value and developing methods to determining a close to optimal value for *ω* is of practical importance and remains to be studied as an independent topic.

It must be noted that scaling the centroid for distance matrix does not follow the same intuition as for the ancestry matrix and cannot be interpreted as strengthening the signals.

## 3 Experimental results

In this section we go through the details of our experimental results. Algorithms and simulations are implemented on Python (3.7.9). ILPs are solved by IBM ILOG CPLEX Optimizer (12.9) with DOcplex (2.23.222) python API.

### 3.1 The synthetic-data generation

To evaluate the average-case performance of the proposed methods on simulated data, first, a set of random labeled rooted trees are generated as the ground truth trees. All generated trees are rooted at 0 depicting germ-line cells. Then, we generate a consistent cancer cell fractions (CCF) randomly in a top-down manner. The CCF value for vertex (mutation) *v* is demonstrated by CCF(*v*). The root vertex is 0 and CCF(0) = 1. For the children of node *v* with specified CCF, the CCFs are chosen uniformly so that their sum is less than or equal to CCF(*v*). This process is repeated until CCFs for all of the vertices are specified. From a single ground truth tree, a set of altered trees are generated by applying one or more of the following alterations (uniformly) at random:

1. **(pc)** (parent-child): This swaps a child vertex with its parent. The chance of this substitution is set to be inversely proportional to the difference between CCFs of the parent and the child vertices.
2. **(bm)** (branch-move): This moves a vertex *v* with all of its descendants to become a child of a vertex *p* with enough *empty CCF* (CCF(*v*) *<* CCF(*p*)− Σ_*u* is child of *p*_ CCF(*u*)).
3. **(nr)** (node-remove): This removes a vertex from where it is to become one of the children of the root. In this process, the vertex former parent becomes the new parent of its former children. The chance of the node-remove alternation for vertex *v* is set to be proportional to 1 − CCF(*v*).

The simulation parameter, **cp** (change-probability), determines the number of changes applied to each tree. Parent-child and branch-move alterations are already applied in simulations reported for ConTreeDP [FS21] and TuELiP [GSO23]. In CeTP, it is assumed that all input trees have the same set of labels (mutations). If some mutations are absent in a number of trees, they can be added as direct children of the root vertex in those trees. This may be considered as the main motivation to add the node-remove alteration to our simulation setup. A realistic scenario in which this alteration (**nr**) seems to be necessary is when multiple bulk samples are taken from the same tumor, while several mutations may not be observable in some of the samples. A detailed explanation of synthetic data generation can be found in Supplementary Section 5.6.

### 3.2 CeTP typically has a unique solution

Note that CeTP does not necessarily admit a unique solution. In this section we study the number of optimal solutions of the optimization problems 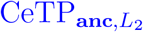, 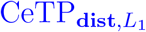, 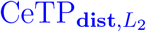 and 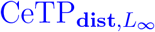. To find the number of optimal solutions, we need to enumerate all trees having the same label set, which is equal to *n*^*n*−2^ by Cayley’s tree formula. In what follows we investigate the case for *n* = 7 vertices where we have reported the results for the subcases with *k* = 3, 10, 15 input trees considering 100 instances of the problem for each *k*.

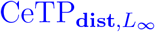 has significantly more solutions per instance than the other three problems (Fig. 3), which seems to be natural since the 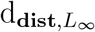 norm has a maximum peak that may allow many small variations in the structures of optimal trees with the same cost. On the other hand, 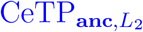, 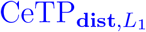 and 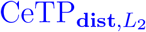 usually have unique solutions, with some sporadic exceptions for which the multiplicity of solutions rarely goes above two. For instance, when *k* = 15, all 100 instances of 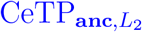 admit unique optimal solutions. Also, more than 50 instances of 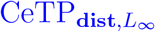 have unique optimal solutions where at least 75 instances have at most 2 optimal solutions (see Fig. 3). It is also interesting to note that the number of optimal solutions decreases as the number of input trees increases (note: 3rd quartiles for all problems are decreasing in Fig. 3).

**Figure 3.**
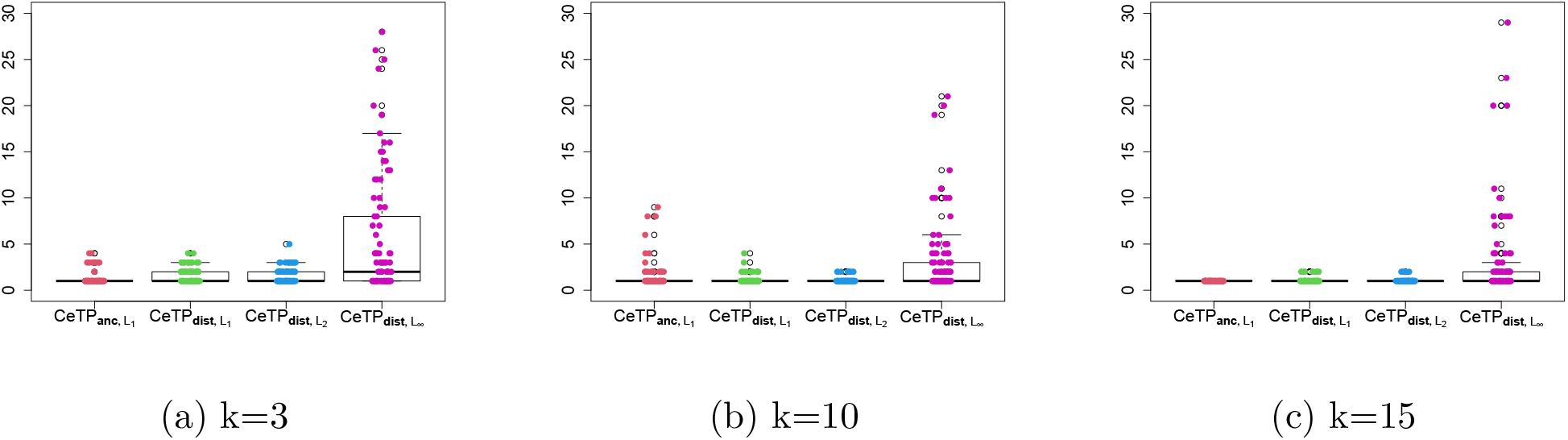
Number of optimal solutions for 100 instances of 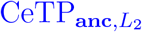, 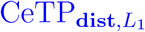, 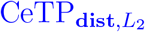, and 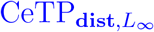 on trees with 7 vertices with *k* = 3 (a), *k* = 10 (b), and *k* = 15 (c) input trees. The *y*-axis shows the number of optimal solutions for instances of these problems and the *x*-axis shows the problems themselves. The number of instances for each simulation setting is set to be 100.

### 3.3 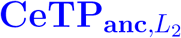 solutions lose most of the ancestry signals

When solving 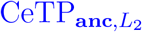 with WAncILP_1_, one observes that many optimal solution trees are star-shaped, indicating that all ancestry relations are lost except for the germ-line cell, which is the ancestor of all vertices in all trees by construction. On the other hand, by increasing the weight of the ancestry relations, it is expected to have a decrease in the number of star-shaped solutions. For our 300 simulations on trees with 7 vertices and *k* = 3, 10, 15, there are 233, 34, and 5 star-shaped trees for solutions of WAncILP_1_, WAncILP_2_, and WAncILP_3_, respectively.

### 3.4 Node-remove alterations produce harder instances to solve

We observed the fact that node-remove alterations add substantial complexity to the problem of finding the ground truth tree. For instance, considering 100 simulations (with parameters *k* = 20, *n* = 10, **cp** = 0.9, **pc** = 1, **bm** = 1), GraPhyC finds 35 true trees when no node-remove change has been applied in the dataset; but when node-remove is added to simulations, one observes that it cannot recover even a single ground truth tree (see Fig. 4) within this dataset.

**Figure 4.**
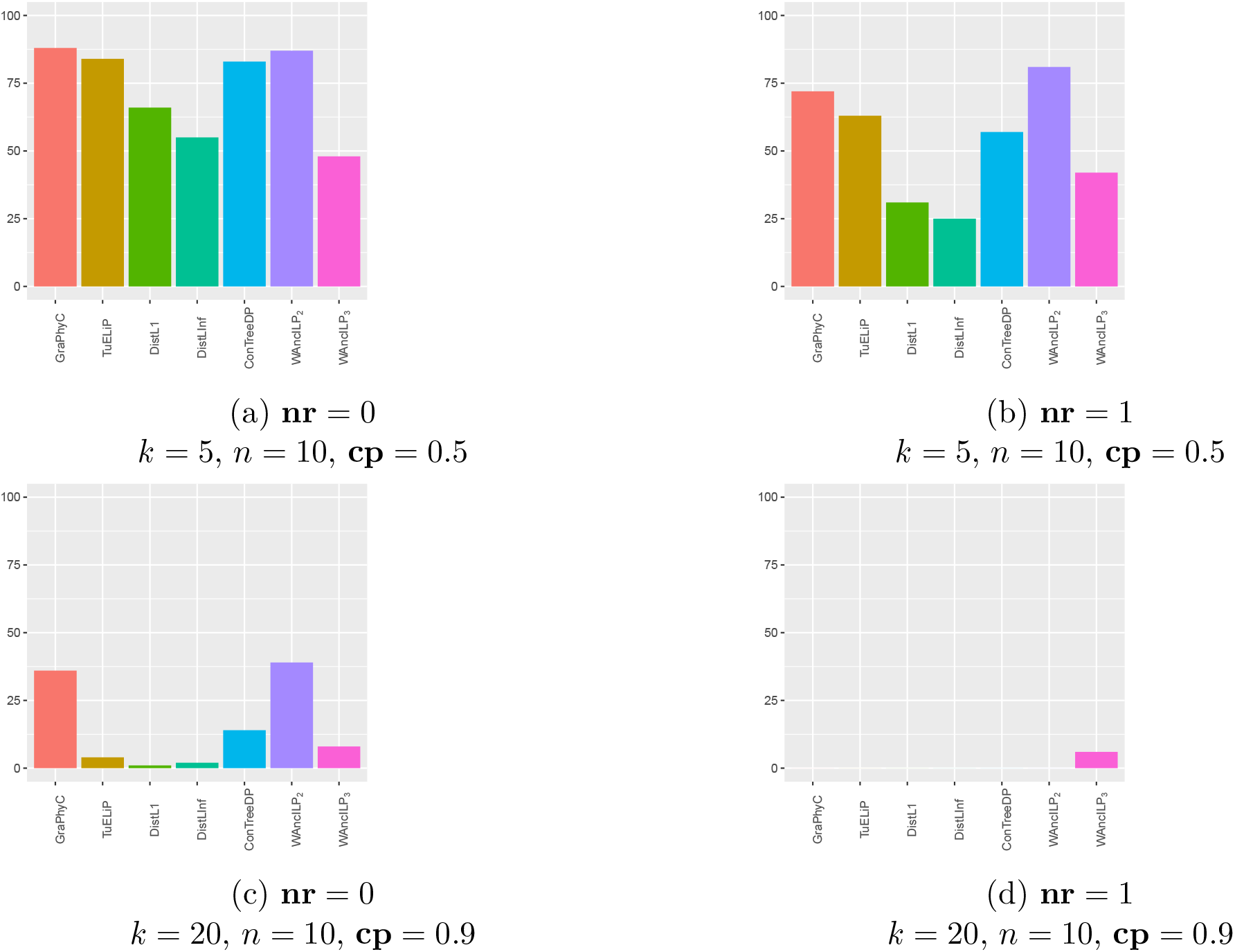
The effect of node-remove alteration in finding the ground truth tree. The plots show the number of trials in which different algorithms output the ground truth tree with respect to, including or excluding, node-remove alterations in the simulation process for two simple simulation settings (a, b) and two hard ones (c, d). Total number of simulations in each setting is 100. Other simulation parameters are **pc** = 1, **bm** = 1.

### 3.5 WAncILP_2_ is a pioneer

We evaluate the algorithms by comparing the distance between their outputs and the ground truth tree for the input set. We use *n* = 10, 20 vertices and 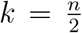, *n*, 2*n* input trees for simulations, with at least one of the three alteration methods (**pc, bm**, and **nr**) active in each simulation setting. This produces 2 × 3 × 7 × simulation settings, with 100 tree sets generated for each setting.

The algorithms are executed on an HPC with 20 cores and 8GB RAM dedicated to each instance of the problem, with a time limit of 4 hours per job. ILPs for **dist** embedding have more variables and conditions than ILPs for **anc**, and consequently, DistL1 and DistLInf are not solved for *n* = 20 within the time limit.

We have used CASet, DISC, PCD, ADD, 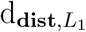, 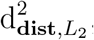, and 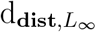 dissimilarity measures to compare the distance between algorithm outputs and the ground truth tree. To evaluate our algorithms (DistL1, DistLInf, WAncILP_2*w*_, and WAncILP_3_), we compare the results with three state-of-the-art algorithms: 1) GraPhyC, 2) TuELiP, and 3) ConTreeDP. Note that TuELiP is applied in its uniformly weighted version because there is no evidence for nonuniform weighting of input trees in our simulation process.

All algorithms behave similarly according to all distance measures except PCD and 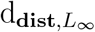 (Supplementary Fig. 14). We have also chosen the CASet distance, already used in evaluations of ConTreeDP and TuELiP, as our measure of comparisons, which is defined as the size of the symmetric difference between the sets of common ancestors for all pairs of mutations in each tree. This, we believe, will set unified framework to be able to provide a fair comparison of performance measures between the prposed algorithms.

**Figure 5.**
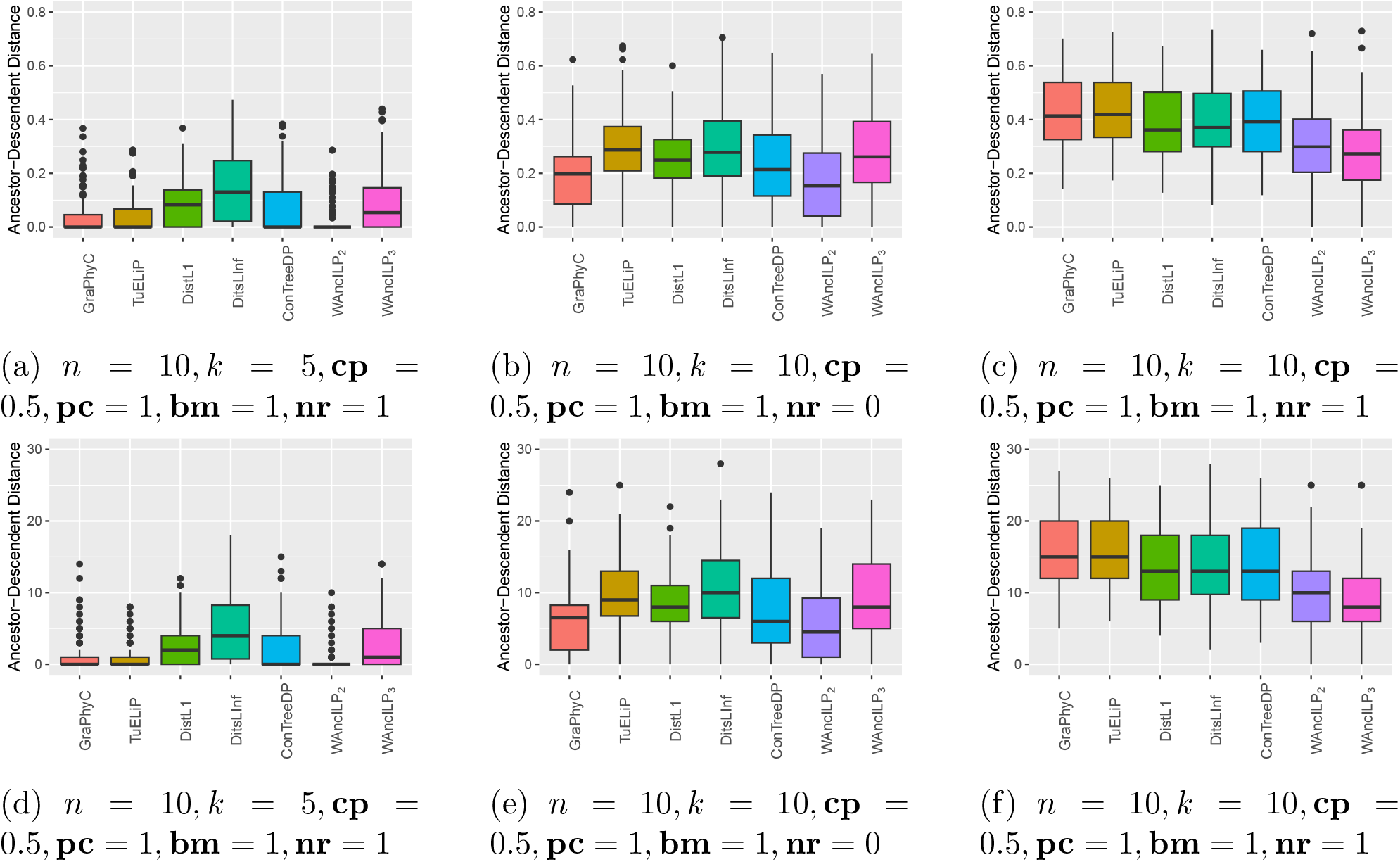
The CASet and ancestor-descendent distance between output of the algorithms and the ground truth tree for three different simulation settings. The *x*-axis shows different algorithms and the *y*-axis shows the distance between output of the algorithm and the ground truth tree.

**Figure 6.**
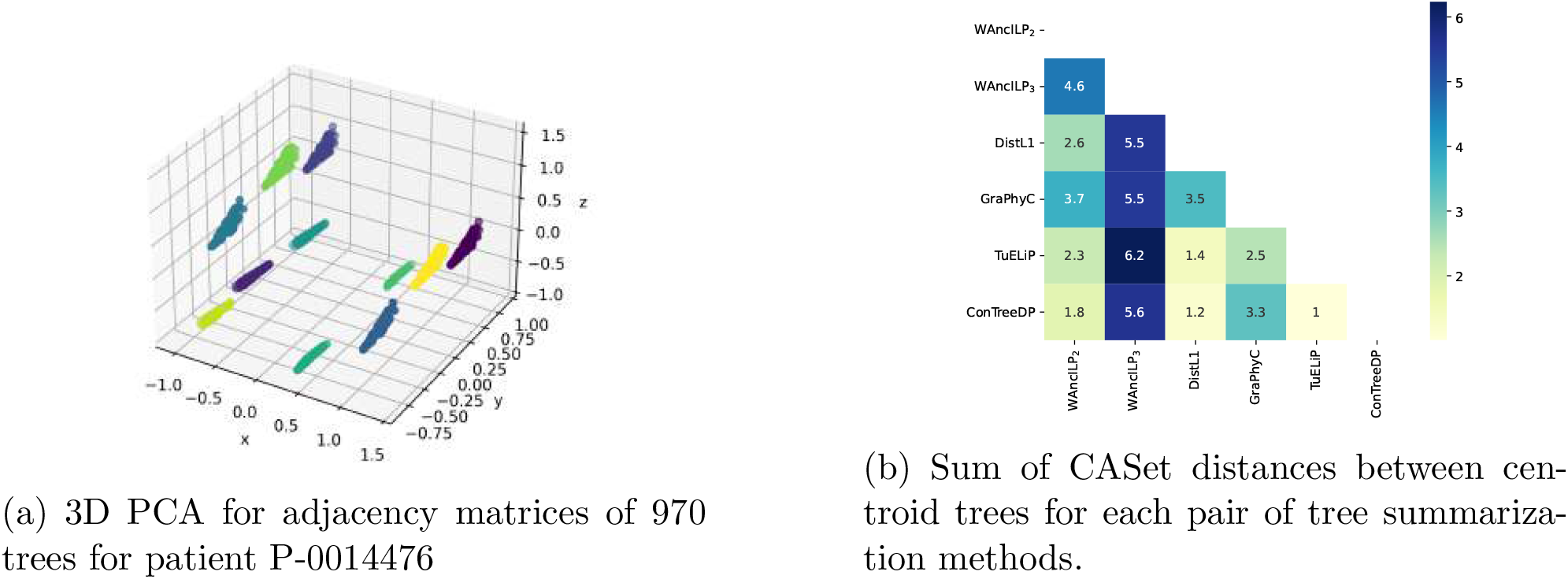
Clustering of trees for patient P-0014476

**Figure 7.**
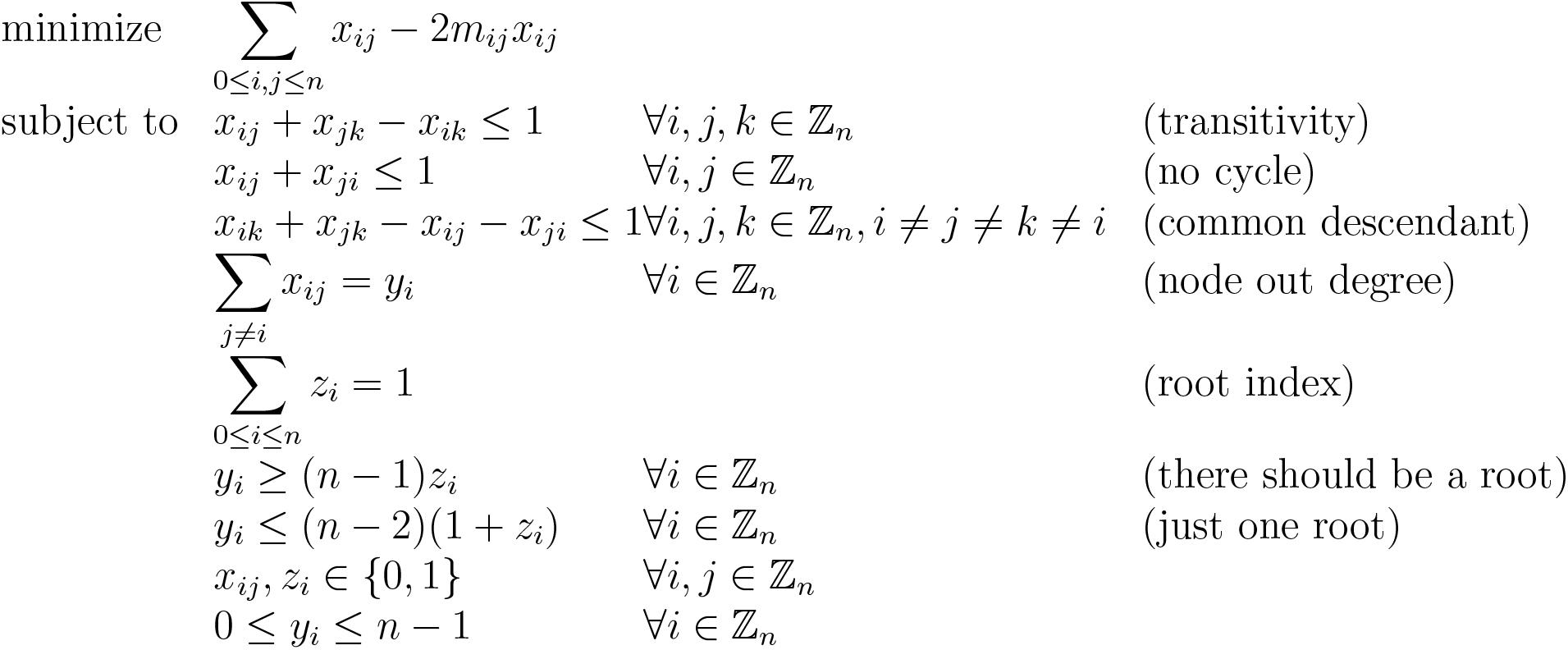
AncL2

**Figure 8.**
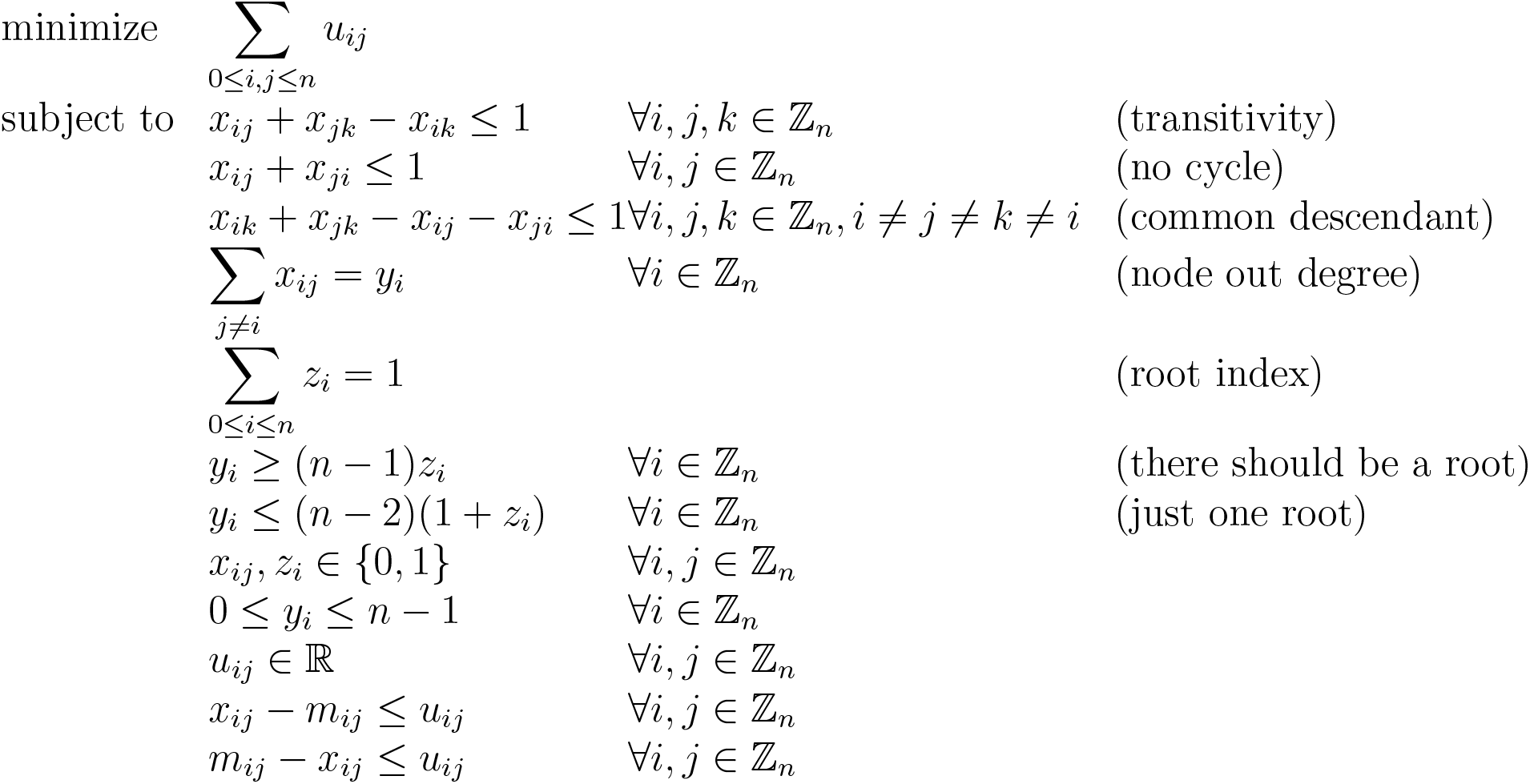
AncL1

**Figure 9.**
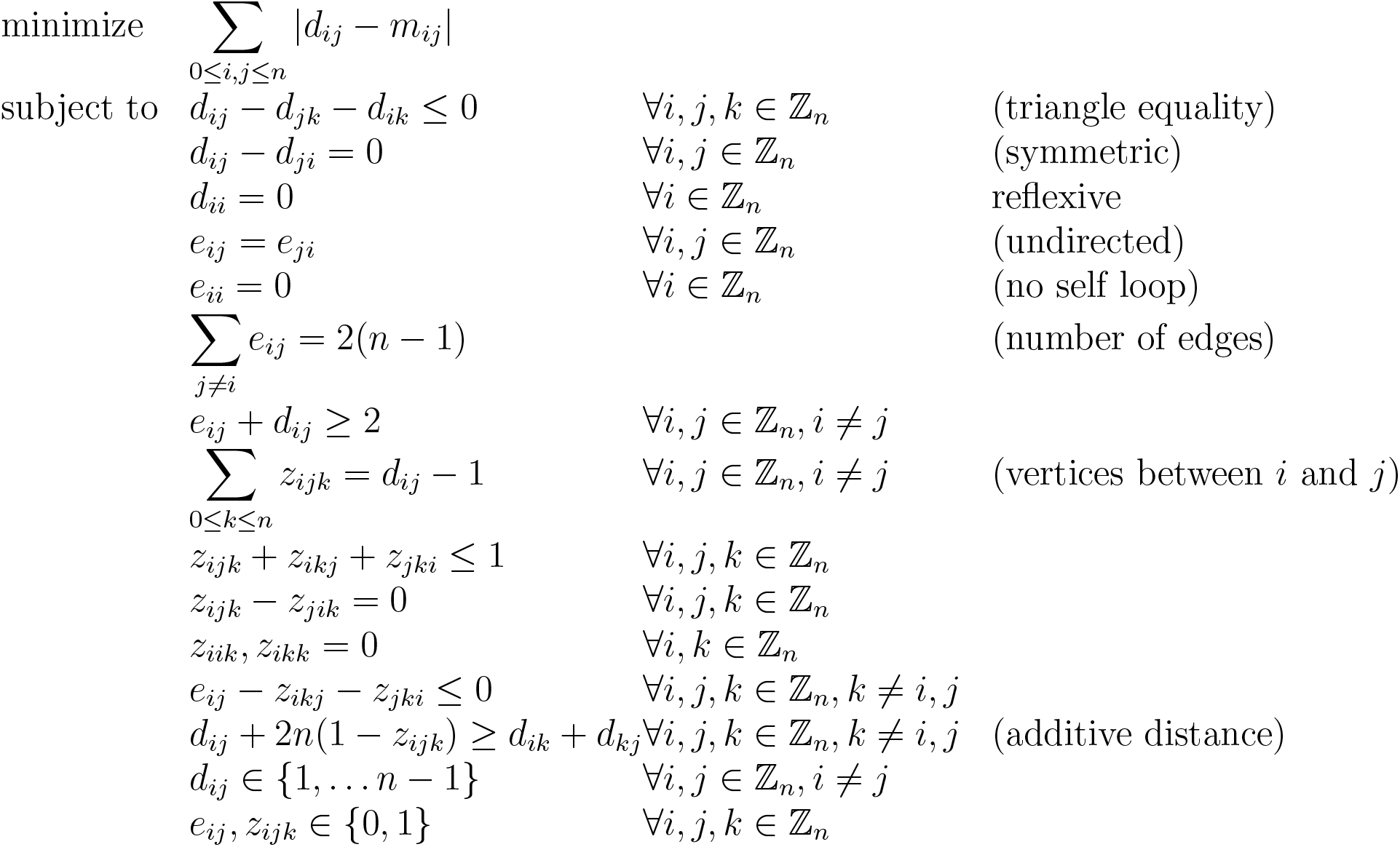
DistL1

**Figure 10.**
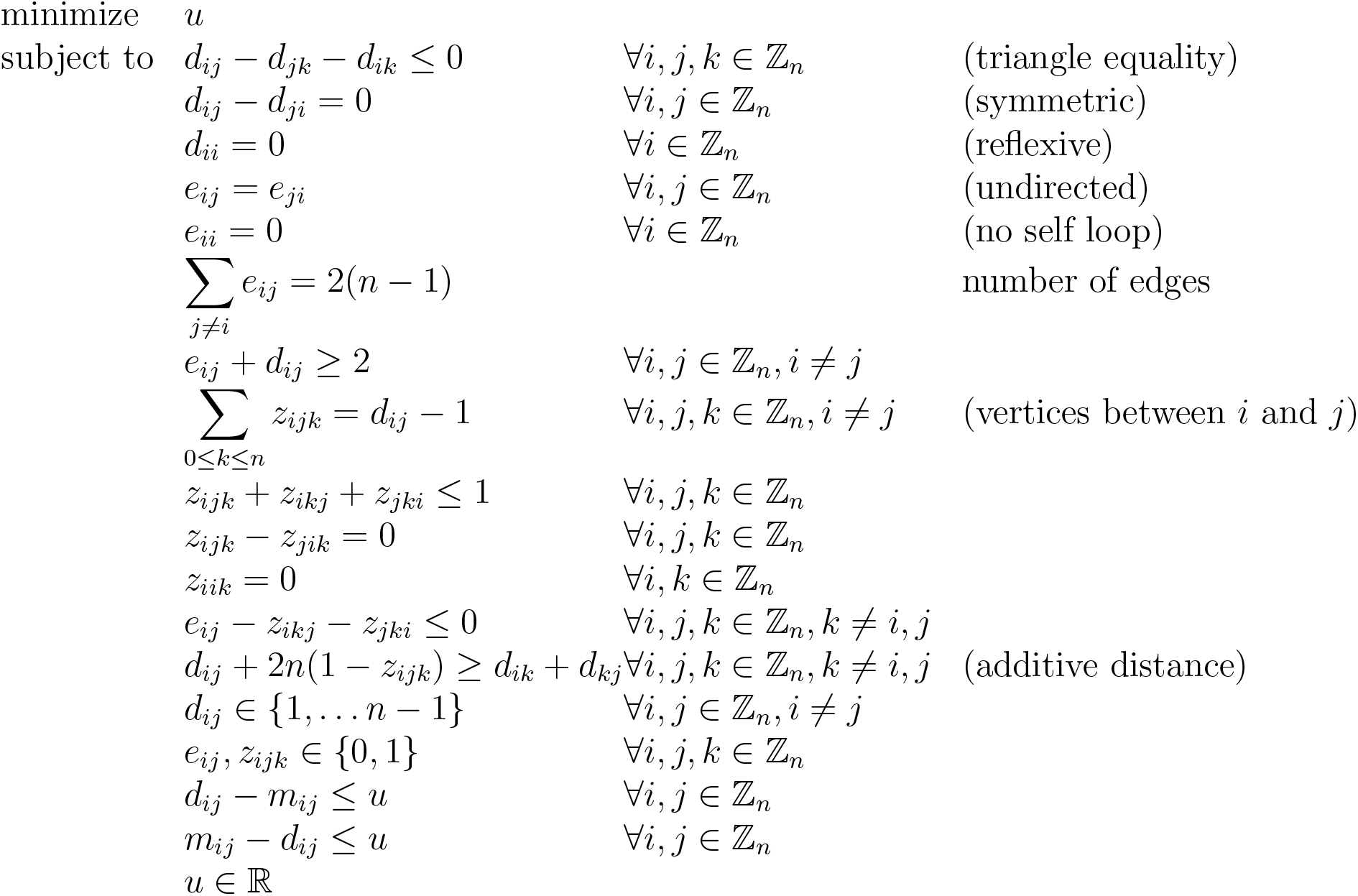
DistLInf

One may observe that WAncILP_2_ is always the best or in the worst case the second-best algorithm in all simulation settings (see Table 2, Fig. 5, and Supplementary Figs. 15 and 16). When the **nr** alteration is present, WAncILP_3_ performs better than all other algorithms, with WAncILP_2_ being the second-best. In the absence of **nr** alterations, WAncILP_2_ and GraPhyC are two of the best-performing algorithms. Also, WAncILP_2_ is better than TuELiP in all simulation settings except when only **pc** alterations are applied (Supplementary Table 4). This in our opinion, is a concrete evidence that the idea of using the weighted setup for the ancestry signal improves the performance of TuELiP’s in WAncILP_2_.

**Table 1:** A list of algorithms and complexity results for different settings of CoTP, CeTP and NMTP problems.

| $\varepsilon$ | $L$ | **CoTP**<sub>$d_{\varepsilon,L}$</sub> | **CeTP**<sub>$\varepsilon,L$</sub> | **NMTP**<sub>$\varepsilon,L$</sub> |
| --- | --- | --- | --- | --- |
| **adj** | $L_1$ | GraPhyC [Gov+20] algorithm<br>$\mathcal{P}$ [Gov+20] | GraPhyC [Gov+20] algorithm<br>$\mathcal{P}$ [Gov+20]<br>(Corollary 1 (a)) | (Supplementary Algorithm)<br>**TrimArb**<br>$\mathcal{P}$<br>(Theorem 6 (b)) |
| **adj** | $L_2$ | GraPhyC [Gov+20] algorithm<br>$\mathcal{P}$ [Gov+20] | GraPhyC [Gov+20] algorithm<br>$\mathcal{P}$ [Gov+20]<br>(Corollary 1 (a)) | Maximum arborescence<br>$\mathcal{P}$<br>(Theorem 6 (a)) |
| **anc** | $L_1$ | TuELiP [GSO23]<br>$\mathcal{NP}$-hard [QE23] | TuELiP [GSO23]<br>$\mathcal{NP}$-hard [QE23]<br>(Corollary 1 (b)) | **AncL1**<br>$\mathcal{NP}$-hard<br>(Corollary 2) |
| **anc** | $L_2$ | TuELiP [GSO23]<br>$\mathcal{NP}$-hard [QE23] | TuELiP [GSO23]<br>$\mathcal{NP}$-hard [QE23]<br>(Corollary 1 (b)) | **AncL2**<br>$\mathcal{NP}$-hard<br>(Corollary 2) |
| **dist** | $L_1$ | No algorithm yet | **DistL1**<br>Complexity unknown | **DistL1**<br>$\mathcal{NP}$-hard<br>(Theorem 10) |
| **dist** | $L_2$ | **DistL2**<br>Complexity unknown<br>(Theorem 5 (b)) | **DistL2**<br>Complexity unknown | **DistL2**<br>$\mathcal{NP}$-hard<br>(Theorem 10) |
| **dist** | $L_\infty$ | No algorithm yet | **DistLInf**<br>Complexity unknown | **DistLInf**<br>Complexity unknown |

**Table 2:** Best performing algorithms in each simulation setting. First and second algorithms with smallest average distance to the ground truth trees are reported for different simulation settings. The significance threshold for p-values is 10^−3^. Given ADD distances of the outputs of algorithms to the ground truth tree, for every pair of algorithms, the p-value is calculated using paired t-test. When the normality condition is not satisfied we have used Kolmogorov-Smirnov test. The null hypothesis is: “distances have the same mean”. When there is no statistical evidence for the superiority of one method, all of the best methods are reported. The number before the name of an algorithm, shows the rank of the algorithm based on its performance. All algorithms in a cell are sorted by the mean value of distances to the ground truth tree. Algorithms in bold are introduced in this article. The change probability (**cp**) parameter is set to be equal to 0.9. [^*^] Parent-child, branch-move, and node-remove parameters for simulation method. The numbers 1 and 0 represent *including* or *excluding* the corresponding alteration, respectively.

| pc, bm, nr* | $\frac{k}{n} = \frac{1}{2}$ | $\frac{k}{n} = 1$ | $\frac{k}{n} = 2$ |
| --- | --- | --- | --- |
| 1, 1, 0 | 1-GraPhyC<br>1-**WAncILP**<sub>2</sub><br>2-ConTreeDP | 1-GraPhyC<br>1-**WAncILP**<sub>2</sub><br>2-ConTreeDP | 1-GraPhyC<br>1-**WAncILP**<sub>2</sub><br>2-ConTreeDP |
| 1, 0, 0 | 1-ConTreeDP<br>2-GraPhyC<br>2-TuELiP<br>2-**WAncILP**<sub>2</sub> | 1-GraPhyC<br>1-TuELiP<br>1-ConTreeDP<br>1-**WAncILP**<sub>2</sub> | 1-GraPhyC<br>1-TuELiP<br>1-**WAncILP**<sub>2</sub><br>1-ConTreeDP |
| 0, 1, 0 | 1-GraPhyC<br>2-**WAncILP**<sub>2</sub> | 1-GraPhyC<br>2-**WAncILP**<sub>2</sub> | 1-GraPhyC<br>2-**WAncILP**<sub>2</sub> |
| 0 or 1, 0 or 1, 1 | 1-**WAncILP**<sub>3</sub><br>2-**WAncILP**<sub>2</sub> | 1-**WAncILP**<sub>3</sub><br>2-**WAncILP**<sub>2</sub> | 1-**WAncILP**<sub>3</sub><br>2-**WAncILP**<sub>2</sub> |

**Table 3:** Rand index and adjusted rand index for comparing cluster agreement between clusters of patient P-0000041’s trees using adjacency, ancestry and distance mappings. Near zero adjusted rand index suggests that these clusters does not agree with each other more than any two random clustering of data with this sizes.

|  | **adj vs anc** | **adj vs dist** | **anc vs dist** |
| --- | --- | --- | --- |
| Rand Index | 0.76 | 0.75 | 0.77 |
| Adjusted Rand Index | 0.06 | 0.02 | 0.14 |

**Table 4:**
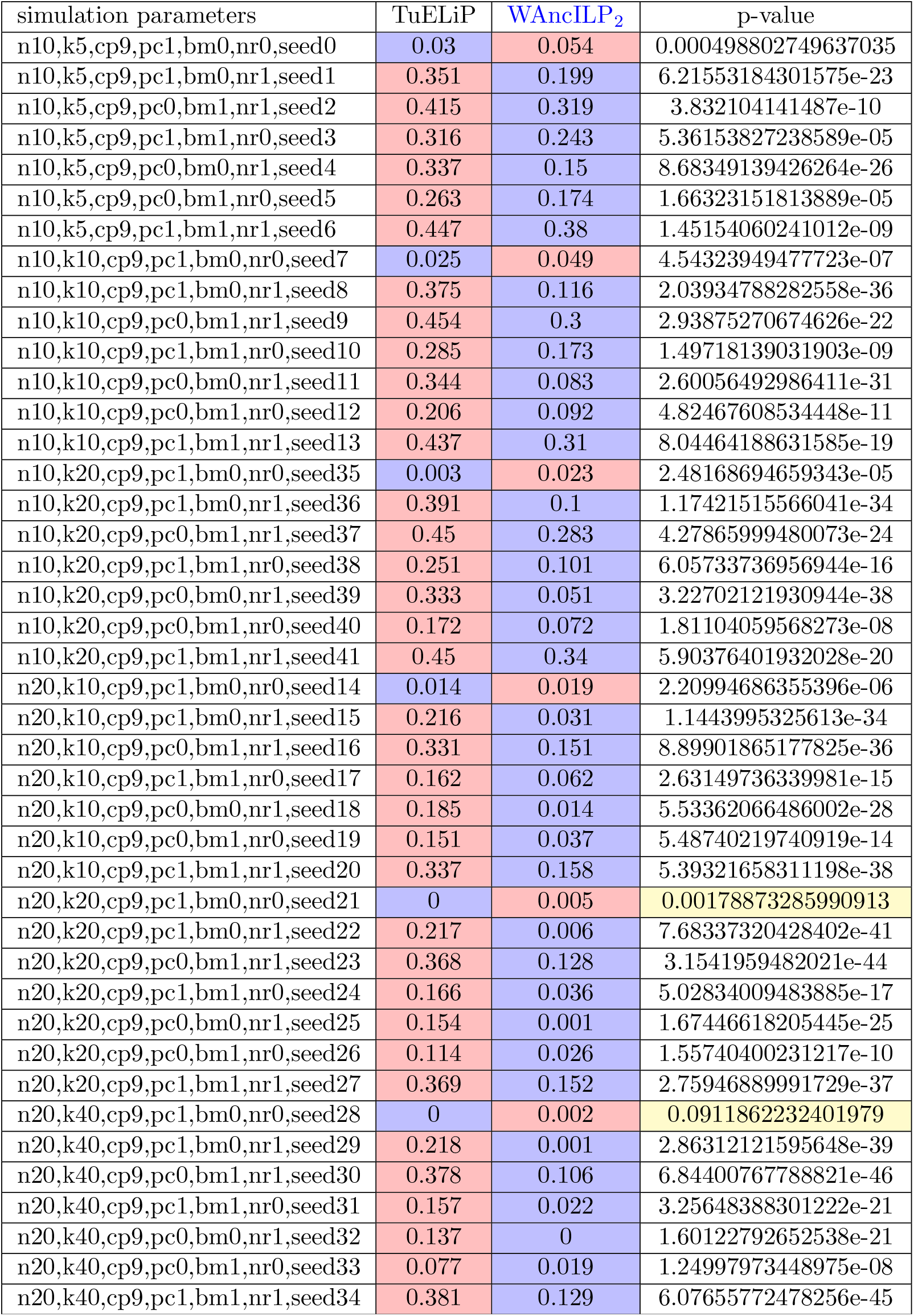
Comparison between mean CASet distance of 100 instances for each simulation settings between TuELiP and WAncILP_2_ algorithms. The p-value is obtained from a paired t-test. The threshold for p-values significance is 10^−3^. The blue color shows smaller mean distance value (red is the opposite). The yellow color specifies insignificant p-values. The dominance of blue value in the WAncILP_2_ column, is an evidence for its better performance.

On the other hand, DistL1 performs quite similar to the ConTreeDP algorithm (Supple-mentary Table 5), where just 5 out of 21 settings show significant difference in the mean distance to the ground truth trees for the two algorithms.

**Table 5:**
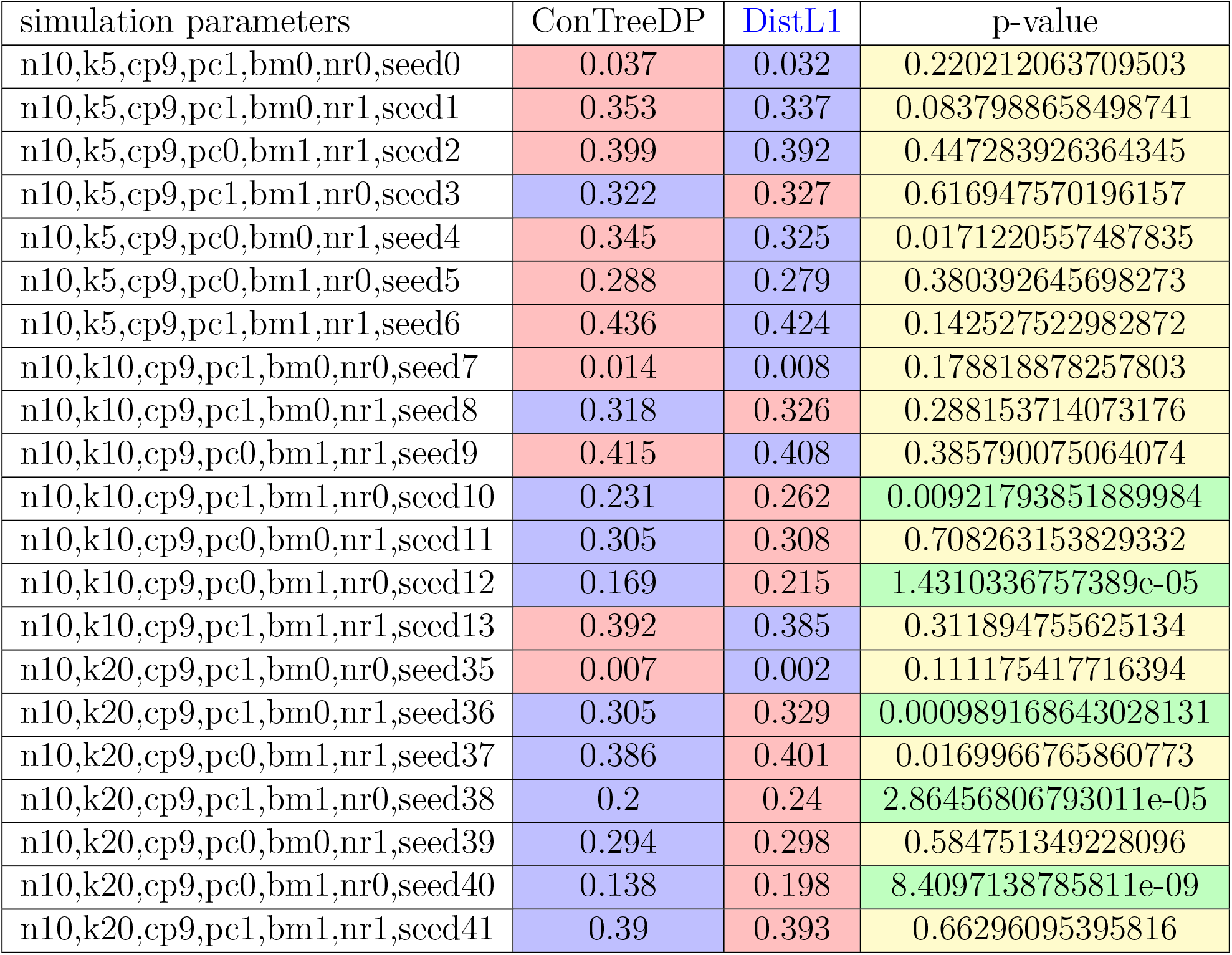
Comparison between mean CASet distance of 100 instances for each simulation settings between ConTreeDP and DistL1 algorithms. The p-value is obtained from a paired t-test. The threshold for p-values significance is 10^−2^. The yellow color specifies insignificant p-values (green is the opposite). The dominance of yellow color suggests that these two algorithms have comparable performance.

We also have noticed that the evaluation process of the algorithms significantly depends on the simulation methods used. For instance, it has already been reported in [FS21] that the ConTreeDP outperforms GraPhyC on almost every simulation set (Supplementary Fig. 12). On the other hand, the dataset used to evaluate TuELiP in [GSO23] shows that GraPhyC and TuELiP outperform ConTreeDP in almost every simulation settings [GSO23]. This sensitivity on simulation strategies has been among our main motivations, not only to provide a general simulation setup covering possible natural alternations along with an exhaustive analysis of different scenarios, but also to generalize the whole problem to the weighted setting that proved to be of importance and use in real applications too (see Section 3.6).

**Figure 11.**
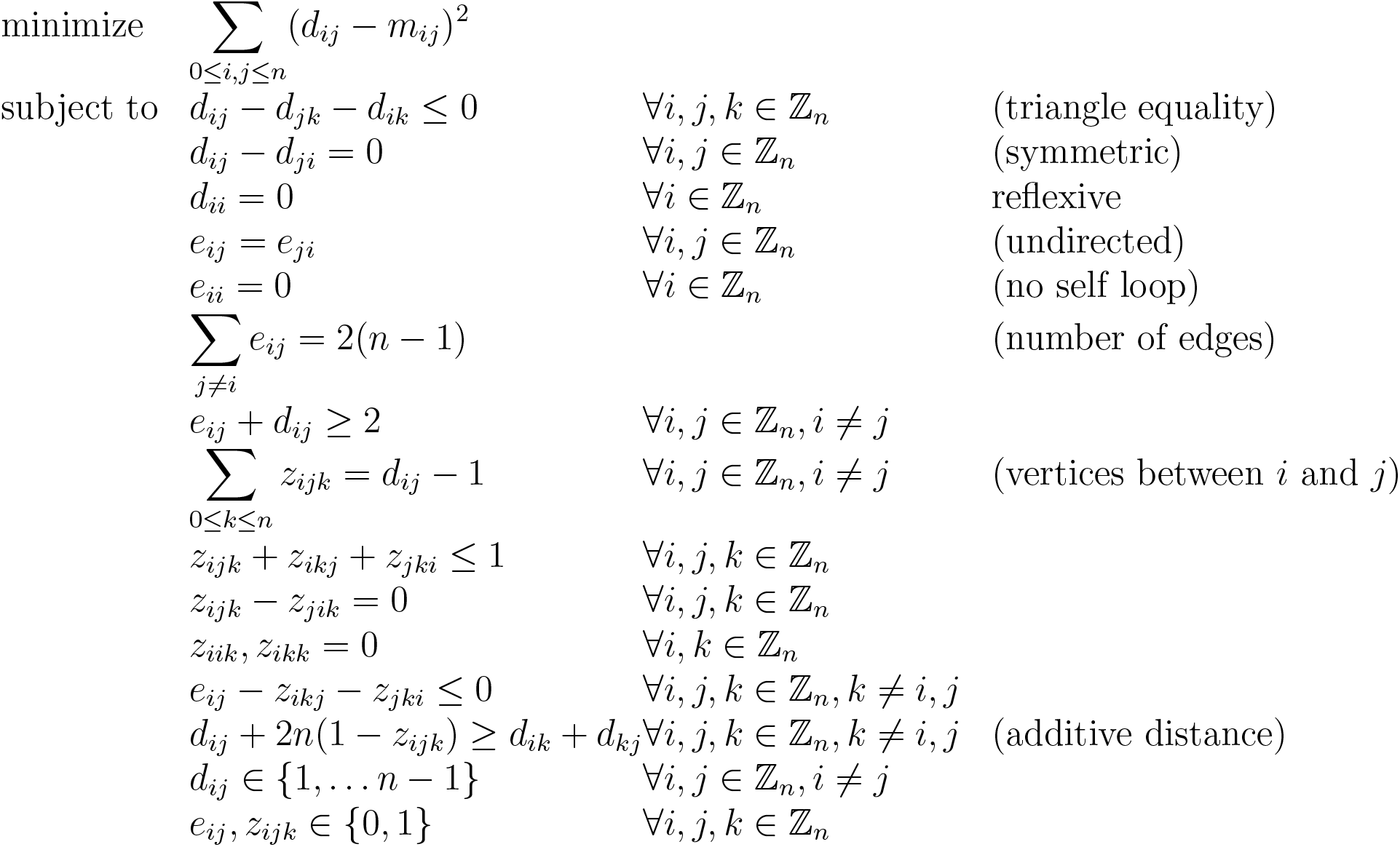
DistL2

**Figure 12.**
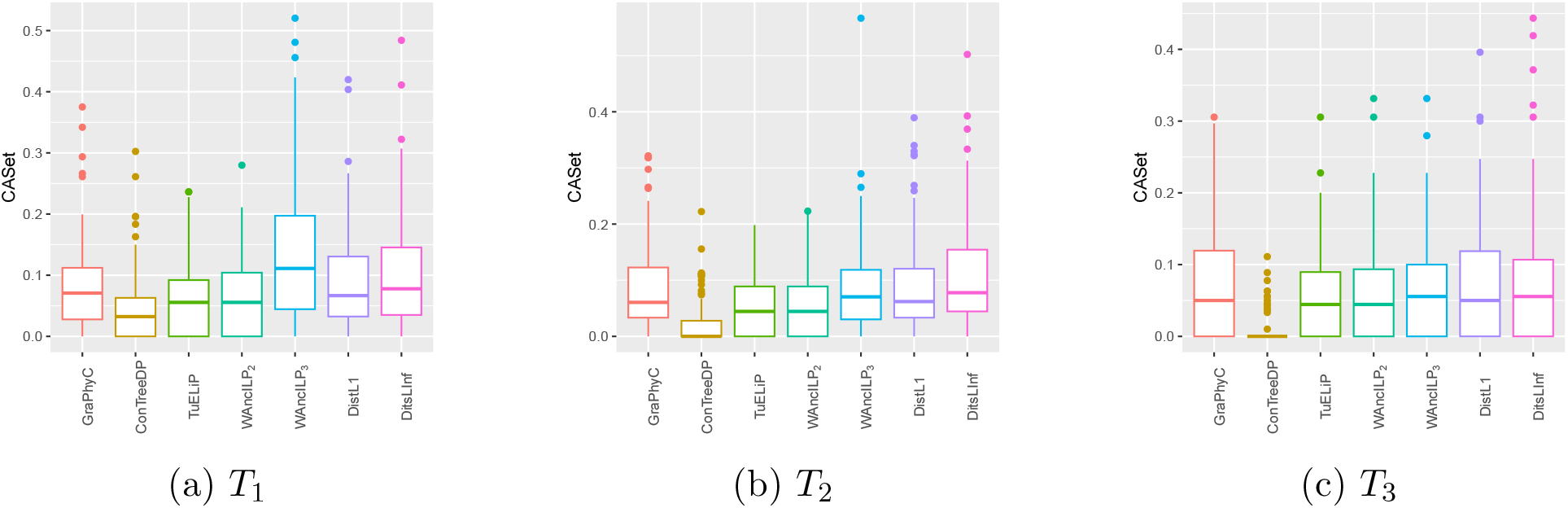
Results of running different algorithms on ConTreeDP study simulated data. As it is obvious ConTreeDP has the best performance on this dataset. It identifies the true structure for most of the trees. It seems very hard to perform better than ConTreeDP on this dataset.

### 3.6 Evaluation on real data

We applied our methods to the real data for breast cancer provided by the cohort study [Raz+18]. The dataset comprises targeted sequencing of 1918 tumors from 1756 patients, focusing on protein-coding exons of 341 to 468 genes that are associated with cancer. We used the tree sets obtained by applying SPRUCE [El-+16] algorithm on this dataset for patients with SNVs occurring in copy neutral autosomal regions. SPRUCE enumerates all tumor phylogenies that explain the variant allele frequencies of the SNVs. These tree sets are available on github page of the RECAP [Chr+20] algorithm.

We picked the patient P-0014476 for our analysis where she has 970 trees with 9 vertices compatible with her data obtained from SPRUCE. Naturally, to study all of these trees, one needs a systematic approach. We, first, applied PCA (principal component analysis) to the adjacency matrices of the trees to reduce the dimensions from 9 to 3, making it possible to visualize all these 970 trees and verify that there are 11 visible clusters for patient P-0014476 in 3 dimensions (see Fig. 13b), certified by applying K-means clustering. The cluster statistics reveal a minimum size of 46, a maximum size of 103, and an average size of 89. In a second stage the algorithms WAncILP_2_, WAncILP_3_, DistL1, TuELiP, GraPhyC and ConTreeDP are applied to obtain a summarization consisting of 11 trees (see Supplementary Figs. 17, 18, 19, 20, 21, 22).

**Figure 13.**
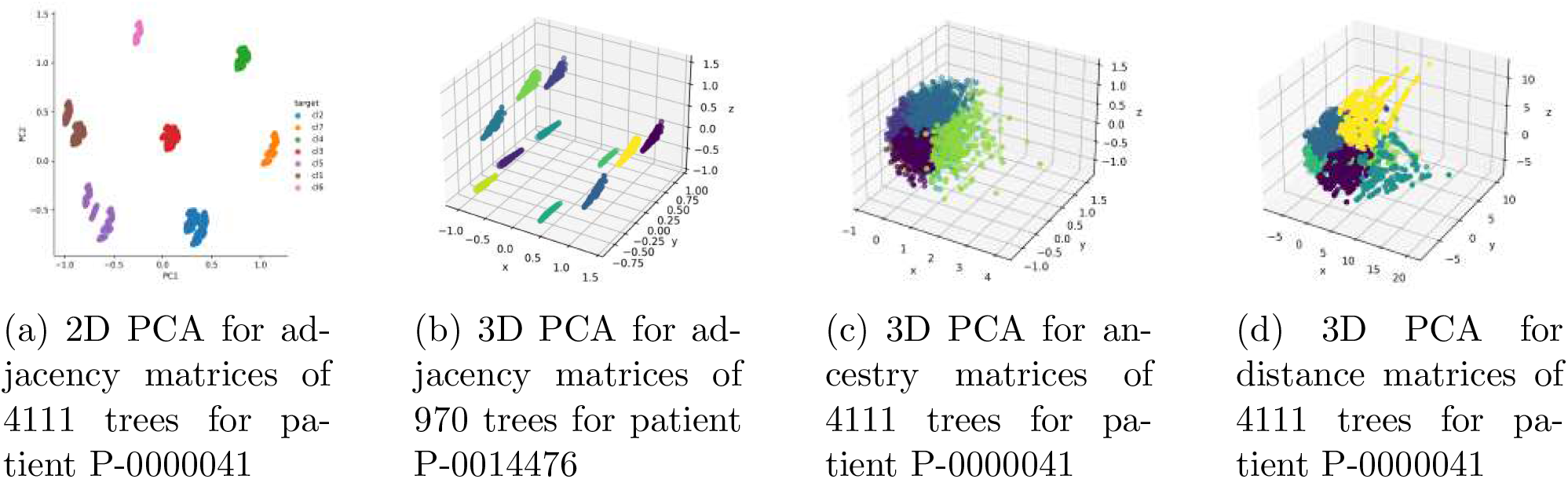
PCA visualization for two patients using **adj, anc**, and **dist** mappings. Near zero adjusted rand index shows low concordance between clusters of different mappings (table 3)

**Figure 14.**
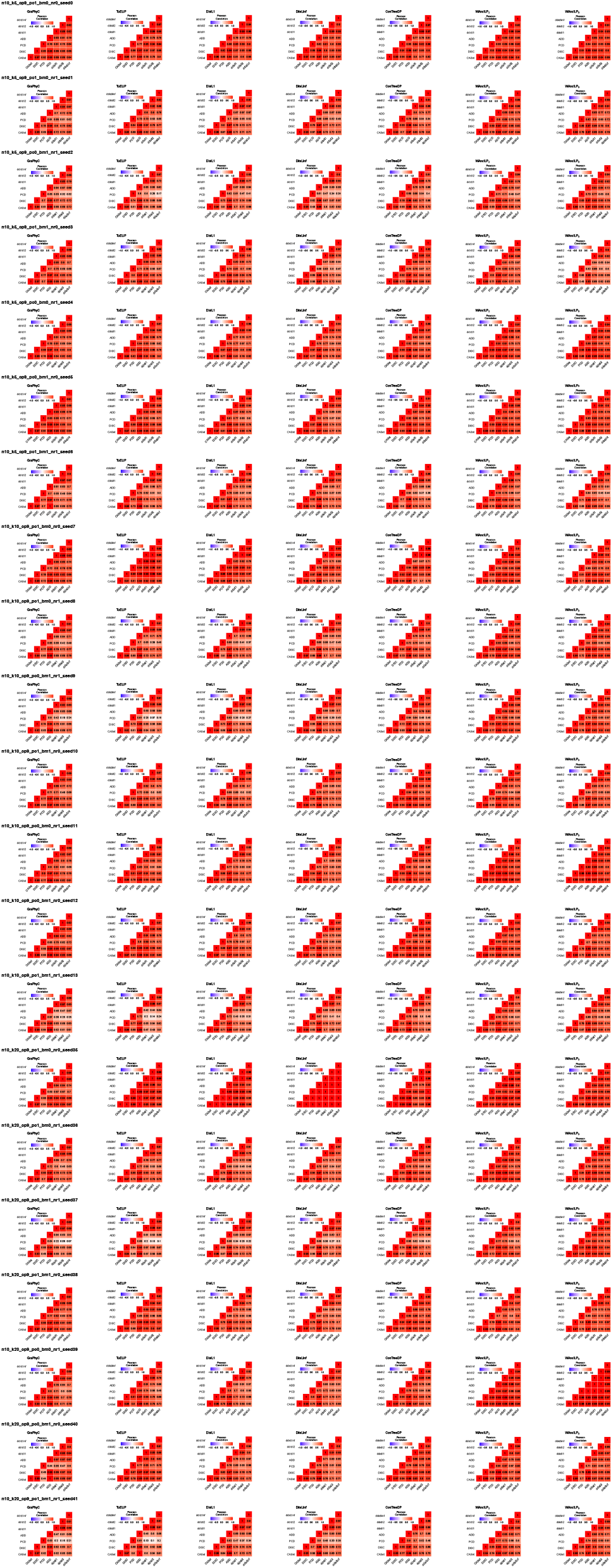
Pearson correlation between different distance measures for solutions of the same algorithm in various simulation settings. Each block shows the Pearson correlation between different distance measures in computing the distance between the solution trees and the ground truth trees for 100 samples of one specific mutation settings solved by a specific algorithm.

**Figure 15.**
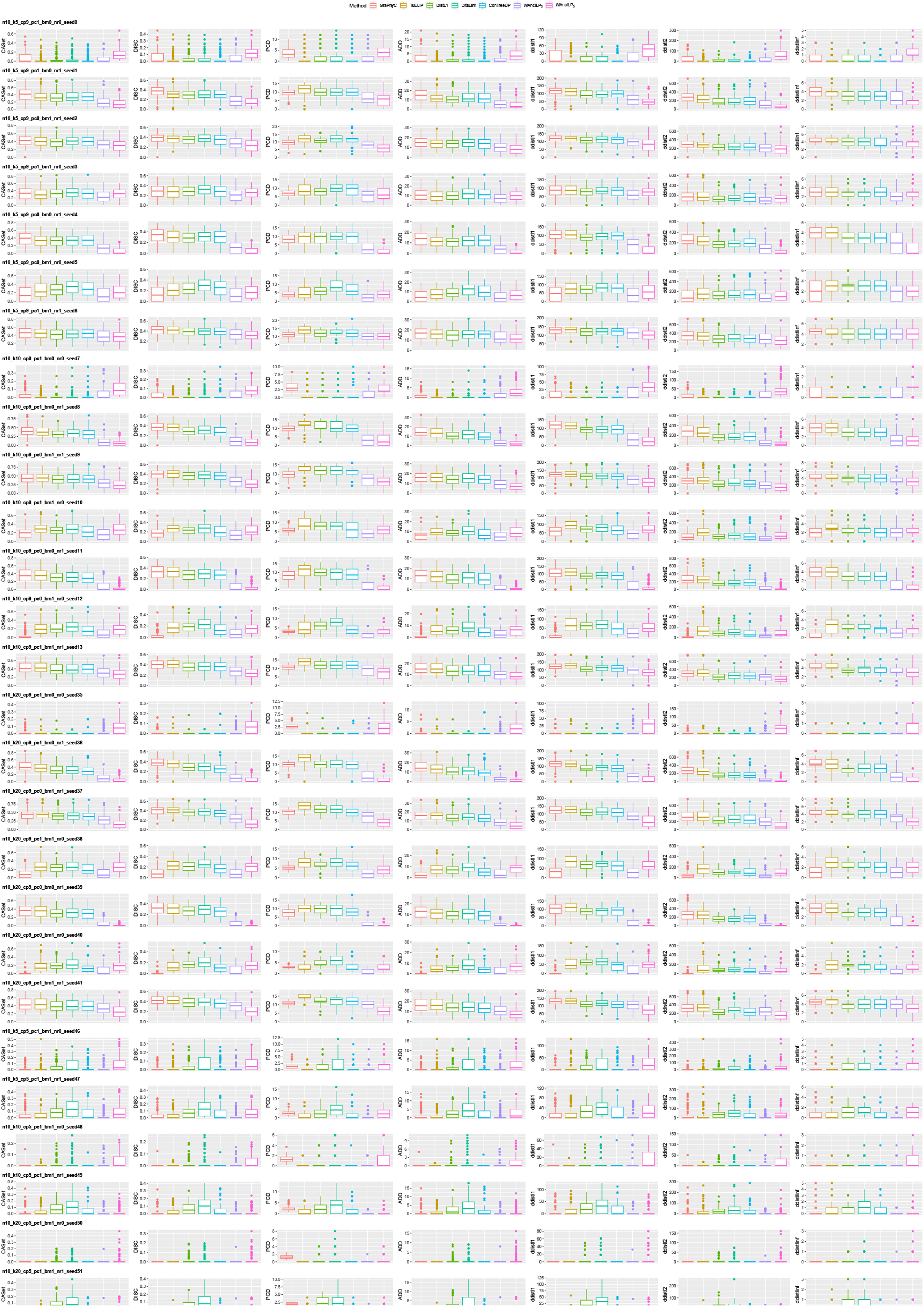
Distance to ground truth tree of outputs of different algorithms in different simulation settings on trees with 10 nodes. A single simulation setting with different distance measures are depicted in each row. Each block shows the box plot of distances between the solutions of different algorithms and the ground truth trees for 100 instances of a specific simulation settings computed by a specific distance measure.

**Figure 16.**
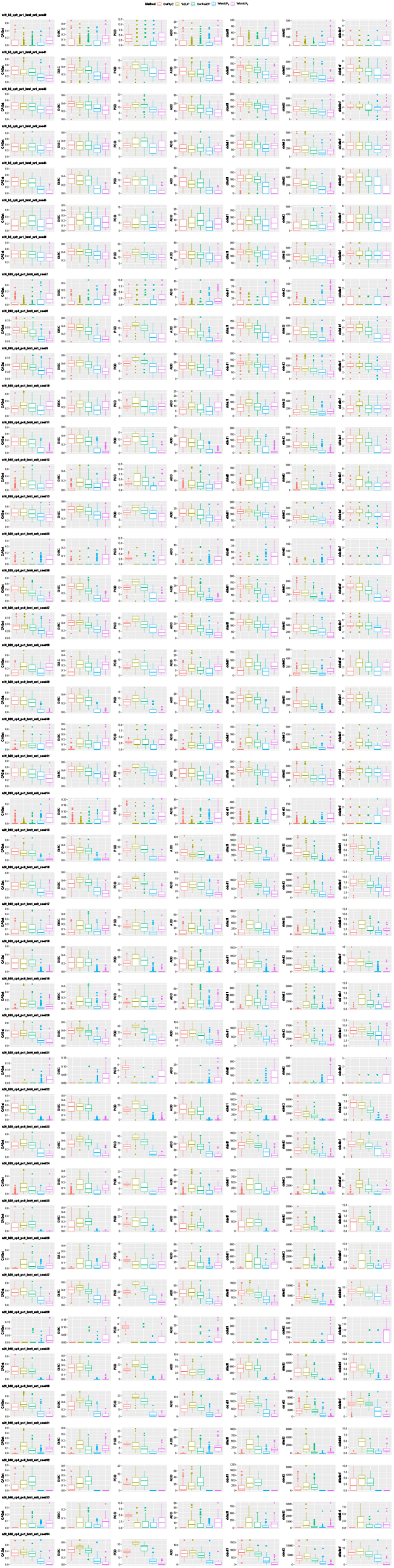
Distance to ground truth tree of outputs of ancestry based algorithms in different simulation settings on trees with 10 or 20 nodes. A single simulation setting with different distance measures are depicted in each row. Each block shows the box plot of distances between the solutions of different algorithms and the ground truth trees for 100 instances of a specific simulation settings computed by a specific distance measure.

**Figure 17.**
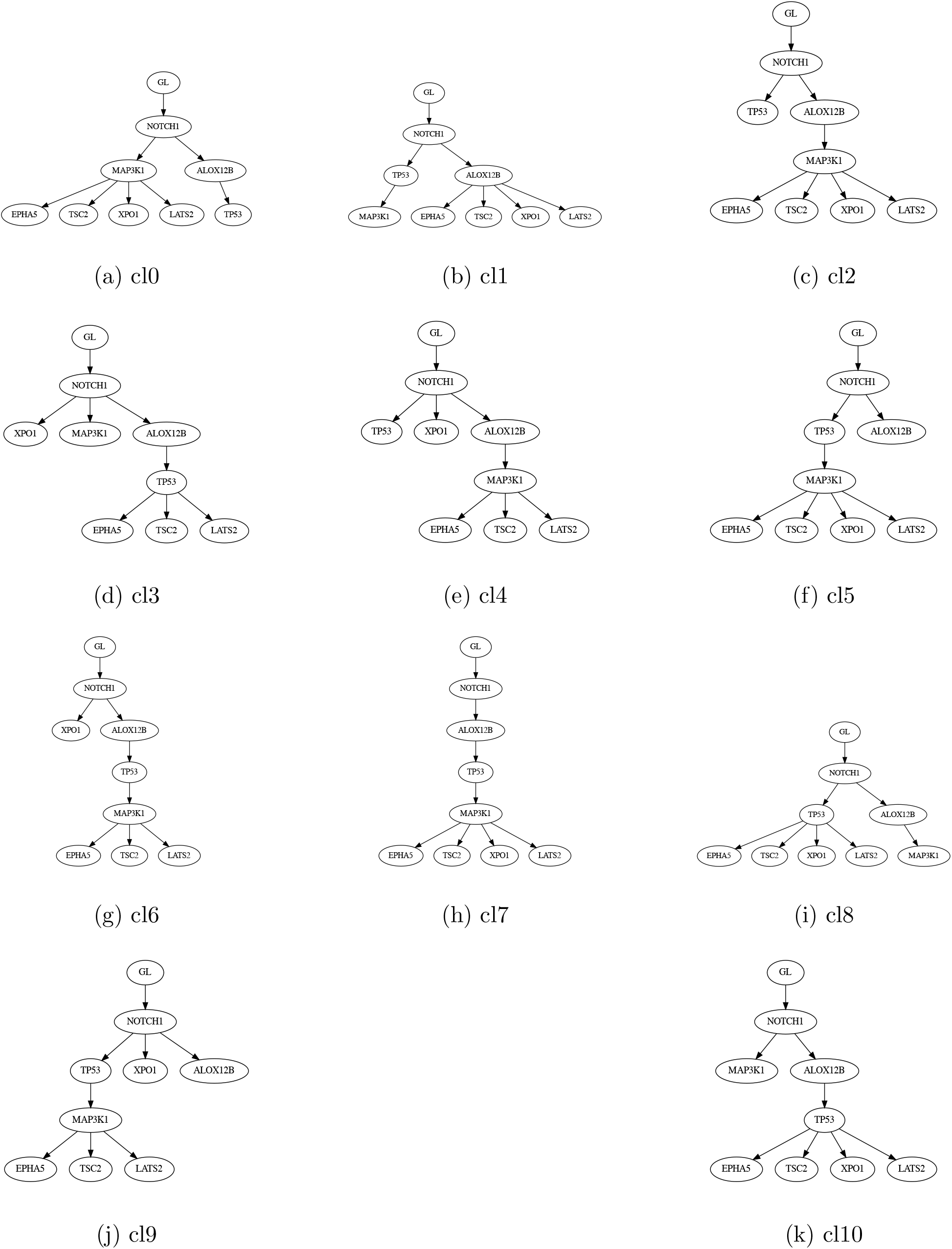
Centroid trees for 11 clusters of adjacency matrices for patient P-0014476 using WAncILP_2_ algorithm.

**Figure 18.**
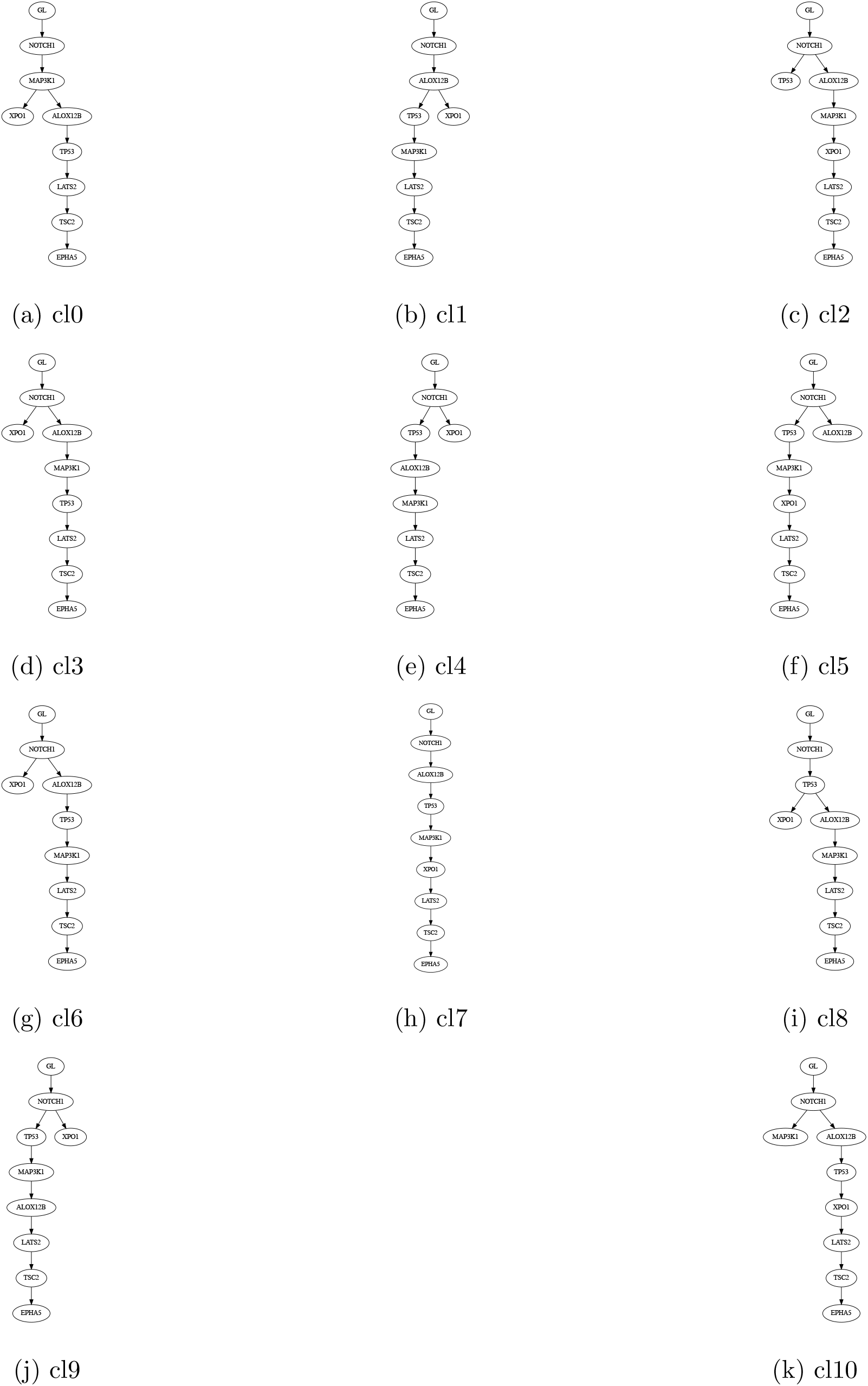
Centroid trees for 11 clusters of adjacency matrices for patient P-0014476 using WAncILP_3_ algorithm.

**Figure 19.**
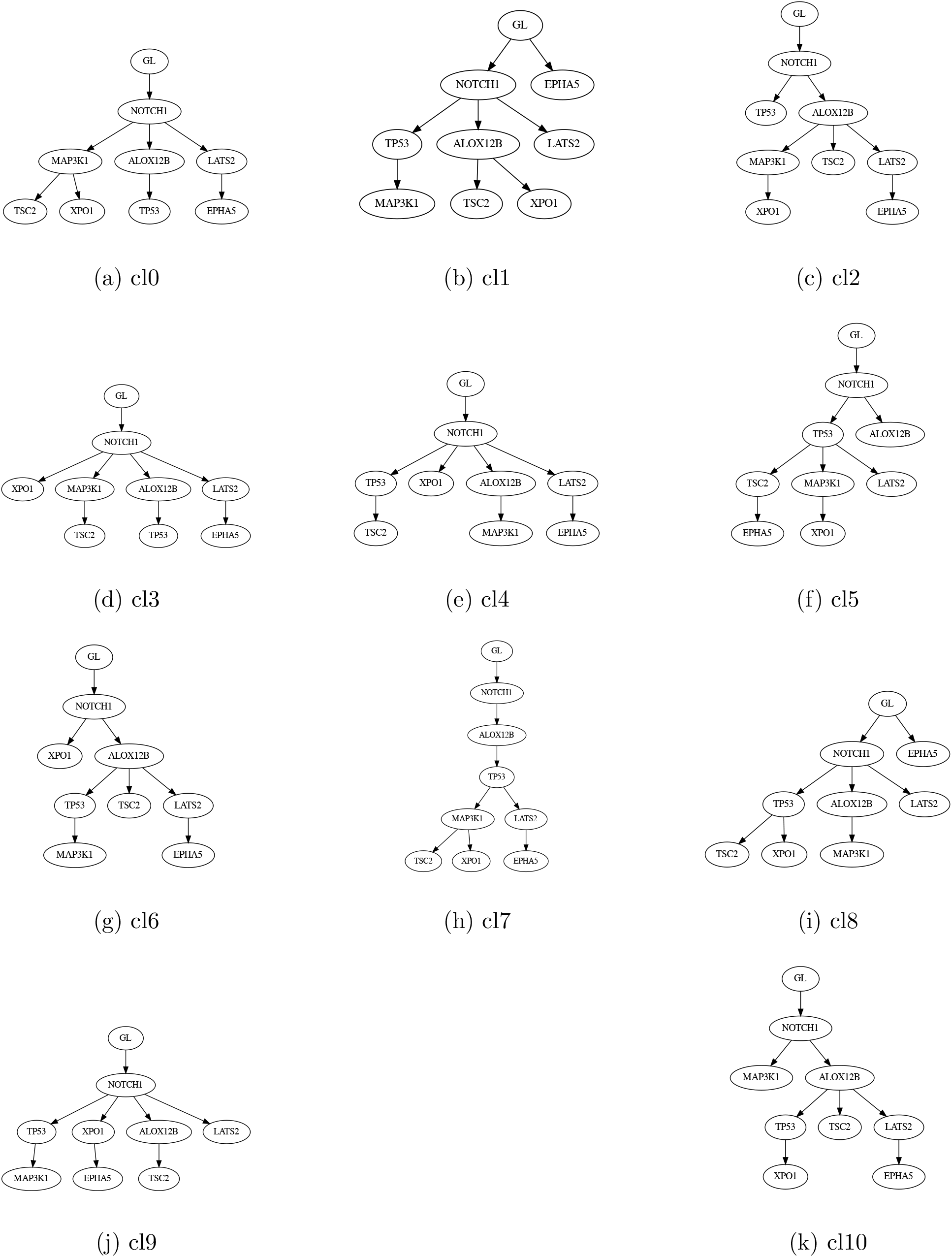
Centroid trees for 11 clusters of adjacency matrices for patient P-0014476 using DistL1 algorithm.

**Figure 20.**
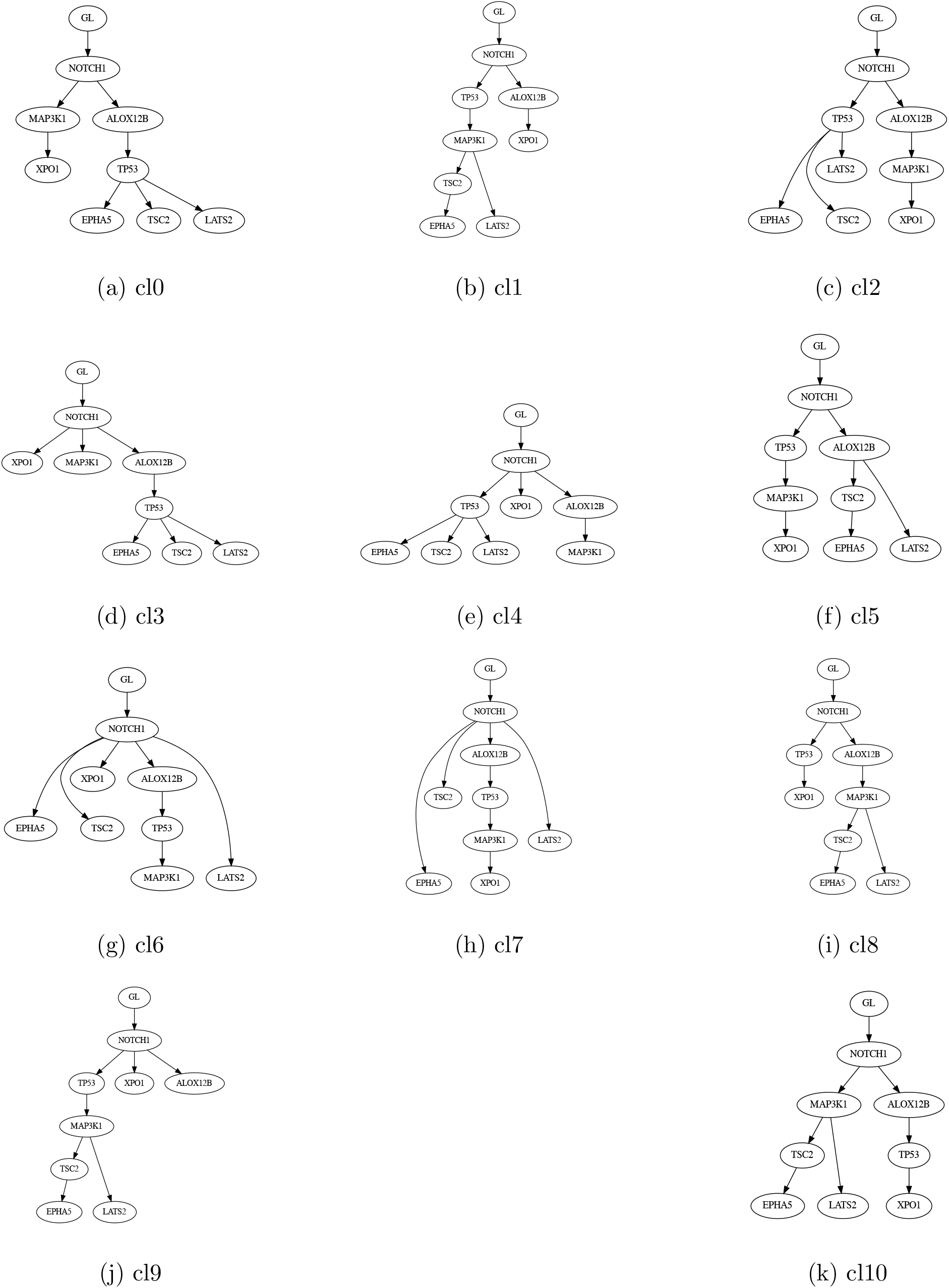
Centroid trees for 11 clusters of adjacency matrices for patient P-0014476 using GraPhyC algorithm.

**Figure 21.**
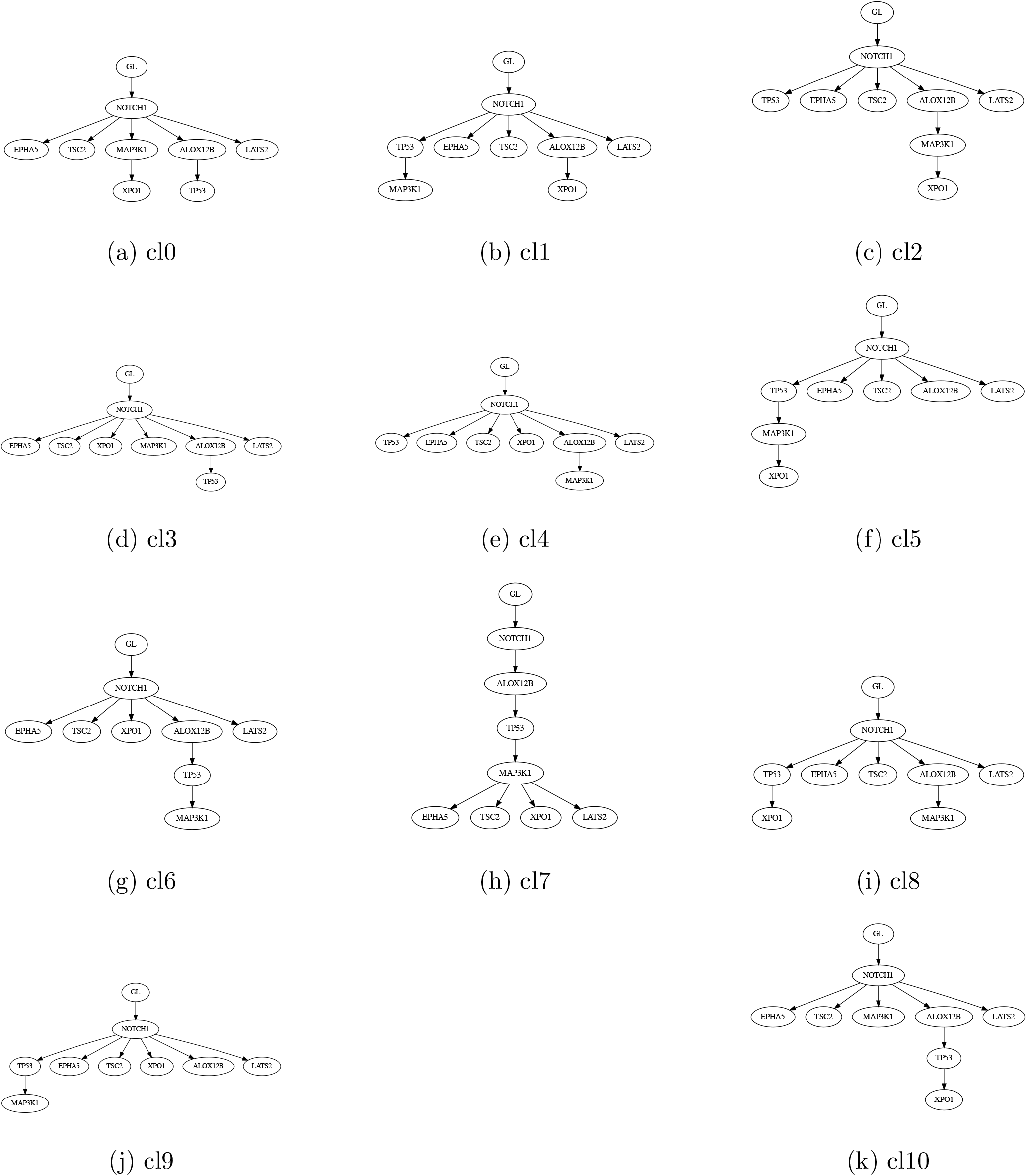
Centroid trees for 11 clusters of adjacency matrices for patient P-0014476 using TuELiP algorithm.

**Figure 22.**
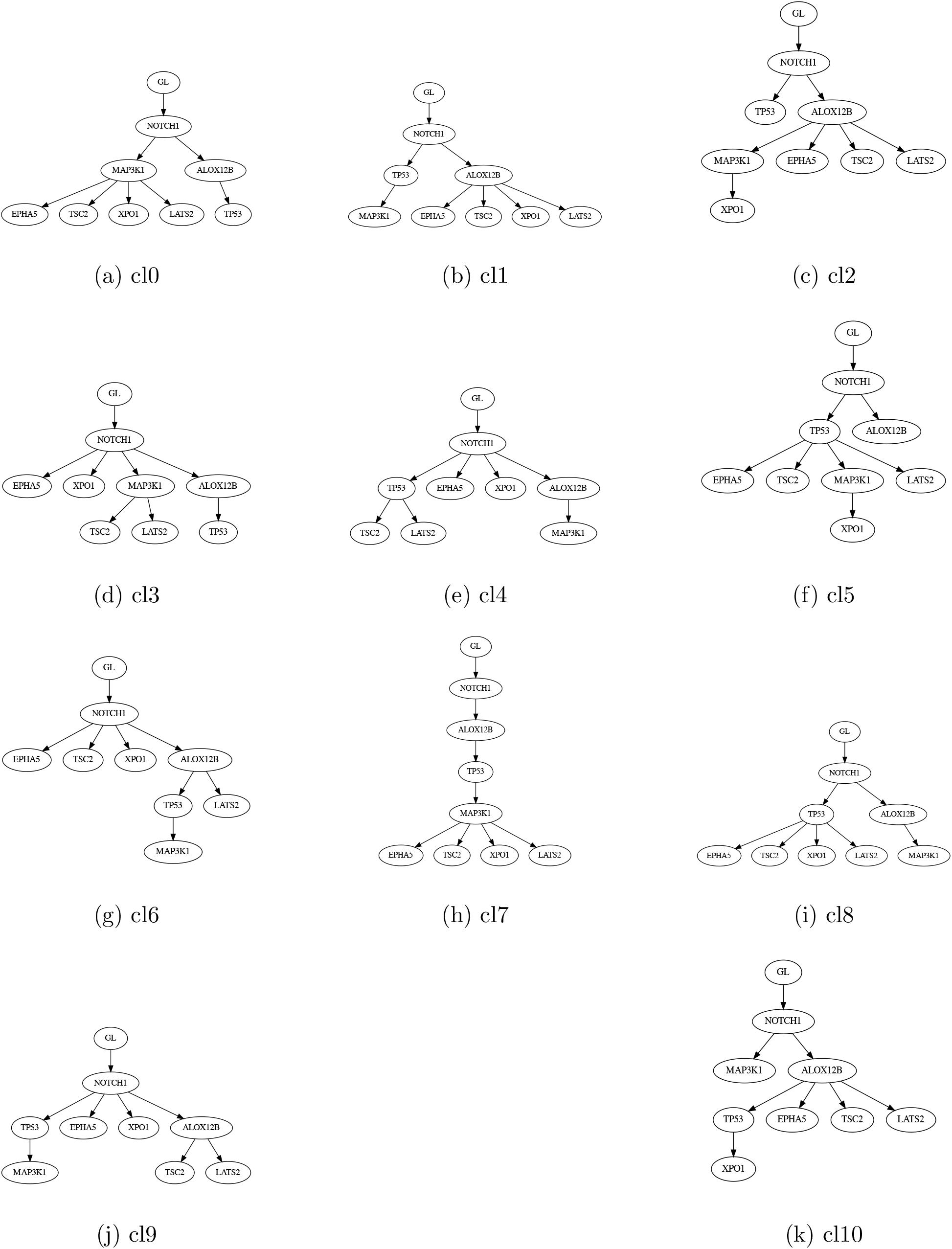
Centroid trees for 11 clusters of adjacency matrices for patient P-0014476 using ConTreeDP algorithm.

The most similar methods are TuELiP and ConTreeDP, however all clusters have different centroid trees obtained from these two methods (Fig. 6b). Also, WAncILP_2_’s centroid trees are similar to ConTreeDP’s, with 4 clusters (cl0, cl1, cl7, cl8) having the same centroid tree for both methods. It ought to be noted that WAncILP_3_ produces the deepest trees, while TuELip produces the shallowest.

Next, we considered centroid trees resulted by WAncILP_2_ algorithm. We observe that NOTCH1 gene is mutated earliest in centroid trees for all clusters using WAncILP_2_ algorithm (Supplementary Fig. 17). The prognostic and regualtory role of Notch signaling pathway in breast cancer has been recently addressed [You+22; Cra+22]. Notch1 signaling encompasses a wide range of functions in determining cell fate, cell proliferation, and apoptosis [Art88; LK06]. Consequently, the NOTCH1 gene has been associated with both oncogenic and tumor suppressor roles [LOA11].

In all centroid trees, NOTCH1 is directly followed by a subset of four genes: ALOX12B, TP53, MAP3K1, and XPO1. This could suggest an early event in carcinogenesis and potentially intimate the causal effects of these mutations in breast cancer. It’s noteworthy that mutations in both TP53 and MAP3K1 have been demonstrated to significantly contribute to the pathogenesis of breast cancer [PAJ13; HG18].

ALOX12B is a gene encoding a lipoxygenase enzyme, and is responsible for the conversion of arachidonic acid to 12R-hydroxyeicosatetraenoic acid [KMF01]. Lipoxygenases are reported to be associated with tumorigenesis and a mutation in ALOX12B has already been detected in lung [She+09] and and cervical cancer [Jia+20]. However, the role of ALOX12B in breast cancer is not well established and limited studies have shown its role in this cancer [Lee+09; Roo+15].

XPO1 is a gene that encodes an export receptor responsible for the nuclear-cytoplasmic transport of hundreds of proteins and multiple RNA species. It is known that this gene is frequently overexpressed and/or mutated in some human cancers (including breast cancer) and plays role as a driver oncogene, making it a good target for breast cancer treatment [AL20; KMS22; Lan+23]. Further research is needed to determine the potential role of ALOX12B and NOTCH1 in development and progression of breast cancer.

## 4 Concluding remarks

We proposed the concept of *weighted centroid trees* to provide a flexible setup for summarizing and analysing classes of mutation trees. Within this framework one is free to choose the embedding map as well as the norm to tune ones best options based on the problem at hand, however, as we have discussed and proved through this contribution, such choices have deep impacts on the solvability of the problem itself. Considering our experimental results, we proposed a new algorithm WAncILP_*ω*_, based on the variant choosing the ancestry matrix for the embedding and the *L*_2_ norm to set the distance, that has a pioneering performance in our experiments among the rest of the algorithms available at the time of writing this article. In this regard we would like to mention some concluding remarks as follows.

It may be verified that some variants of the problem may not be interesting for real applications in phylogenetics, but some particular instances of the problem as 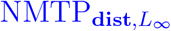 may still be of independent interest, because of the trade-off between the hardness imposed by the **dist** embedding and the simplicity imposed by the maximum norm. Similarly, using the same techniques we used to prove ILPs, one may provide quadratic programs for some other variants of the problem as 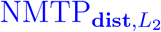, which are not quite efficient in implementation. Needless to say, a study of computational complexity of such variants of the problem and looking for effective approximation algorithms in each case are among the interesting theoretical byproducts of our research.

Although our setup is flexible enough to let weighting strategies for the input trees, as already applied in TuELiP, our generalized set up also makes it possible to apply weighting strategies at a deeper level, to tune the centroids, and consequently, the resulting representative tree at the output. This new weighting strategy clearly reveals its importance when one is facing node-remove alterations as reported in our experimental results. On the other hand, finding efficient methods to determine the best choice for the weighting function in this setup, and in particular for the WAncILP_*ω*_ algorithm, is again among the interesting problems that ought to be studied in future research.

To do our best, we applied our algorithm to both synthetic data and an available real breast cancer dataset. This enabled us to systematically explore a large number of trees and report the effectiveness of our methods in correlation with what is already known in this context, proving that our proposed algorithm has an acceptable performance in real cases as well. However, despite being seemingly comprehensive, this still calls for the development of a reliable and standard benchmarking dataset in this field of research.

## 5 Supplementary Materials

### 5.1 Notations

In addition to notations already defined in the main text, In this text we denote vectors by lowercase boldface letters (like **a**). Also,, the *inner product* of two matrices is defined as

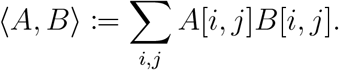

Note that for any real number *r* we have

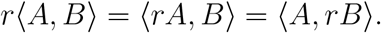

### 5.2 Dissimilarity measures

#### Proposition 1.

The dissimilarity measure d_*ε*,*L*_ is a metric *if and only if ε* : *𝒯*_*n*_ → ℝ ^*n×n*^ is one-to-one.

*Proof*. If *ε* is not one-to-one then there exist two distinct trees *T*_1_ and *T*_2_ for which *ε*(*T*_1_) = *ε*(*T*_2_) hence d_*ε*,*L*_(*T*_1_, *T*_2_) = 0 and d_*ε*,*L*_ is not a metric. Conversely, *if ε* is one-to-one then d_*ε*,*L*_ is a metric:

1. The distance from a point to itself is zero, since d_*ε*,*L*_(*T*_1_, *T*_1_) = *L*(0) = 0.
2. (Positivity) The distance between two distinct points is always positive, since if *T*_1_ ≠ *T*_2_, then *ε*(*T*_1_) ≠ *ε*(*T*_2_), implying *x* = *ε*(*T*_1_) − *ε*(*T*_2_) ≠ 0, and consequently, d_*ε*,*L*_(*T*_1_, *T*_2_) = *L*(*x*) ≠ 0.
3. (Symmetry) The distance from *T*_1_ to *T*_2_ is the same as the distance from *T*_2_ to *T*_1_, since d_*ε*,*L*_(*T*_1_, *T*_2_) = *L*(*ε*(*T*_1_) − *ε*(*T*_2_)) = *L*(*ε*(*T*_2_) − *ε*(*T*_1_)) = d_*ε*,*L*_(*T*_1_, *T*_2_).
4. The triangle inequality holds, since d_*ε*,*L*_(*T*_*x*_, *T*_*z*_) = *L*(*ε*(*T*_*x*_) − *ε*(*T*_*z*_)) = *L*(*ε*(*T*_*x*_) − *ε*(*T*_*y*_) + *ε*(*T*_*y*_) − *ε*(*T*_*z*_)) ≤ *L*(*ε*(*T*_*x*_) − *ε*(*T*_*y*_)) + *L*(*ε*(*T*_*y*_) − *ε*(*T*_*z*_)) = d_*ε*,*L*_(*T*_*x*_, *T*_*y*_) + d_*ε*,*L*_(*T*_*y*_, *T*_*z*_). □

#### Proposition 2.

All dissimilarity measures 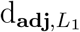, 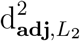, 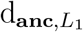, 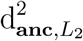, 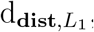, 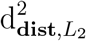 and 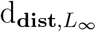 are computable in *𝒪*(*n*^2^) time. □

*Proof*. All three matrices for adjacency, ancestry, and distance can be computed in *𝒪*(*n*^2^) time, by running a BFS from the root towards the leaves and updating *𝒪*(*n*) entries at each round. Comparing two matrices element-wise is also possible in *𝒪*(*n*^2^) time.

#### 5.2.1 PCD and ADD

##### Definition 4 (PCD).

([Gov+20]) The *parent child set* of a tree *T* is defined as

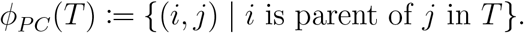

Given two mutation trees *T*_1_ and *T*_2_, the *parent-child distance* is defined as

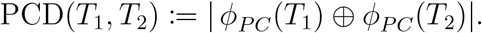

##### Definition 5 (ADD).

([Gov+20]) The *ancestor descendent set* of a tree *T* is defined as

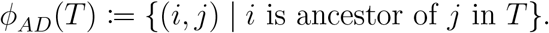

Given two trees *T*_1_ and *T*_2_, the *ancestor-descendent distance* is defined as

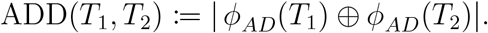

**Observation 5.1**. For two mutation trees *T*_1_, *T*_2_ ∈ *𝒯*_*n*_ we have

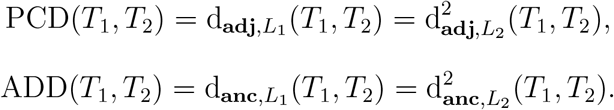

*Proof*. Since the elements of the adjacency matrix are 0 or 1 we have

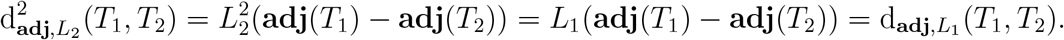

On the other hand,

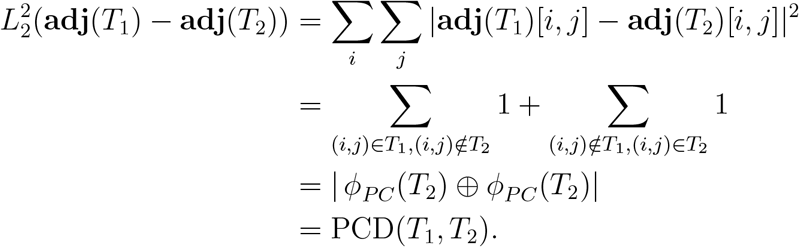

Since the elements of the ancestry matrix are 0 or 1 we have

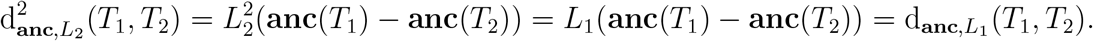

On the other hand,

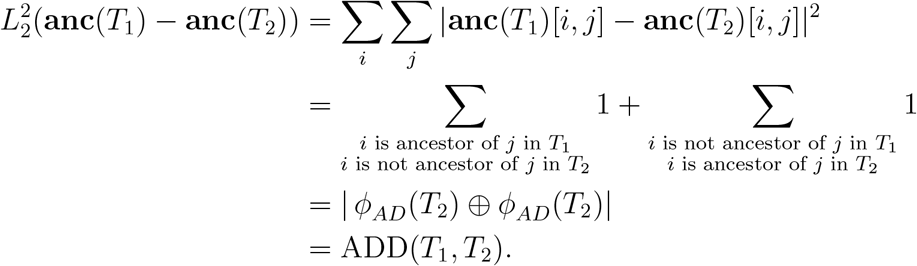

##### Corollary 4.

Observation 5.1 and Proposition 1 imply that 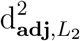 and 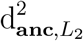 are metrics over trees.

### 5.3 Consensus vs Centroid Tree

#### Lemma 1.

Given a set of matrices S = {**A**_1_, …, **A**_*k*_} whose entries are in [0, 1], a weight vector **w** = {*w*_1_, …, *w*_*k*_} whose entries are also in [0, 1] and for which 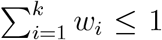, and a real number *r >* 0, the following cost functions have equal minimizers over any subset of binary matrices, i.e., matrices with elements in {0, 1}:

1. 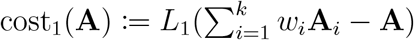,
2. 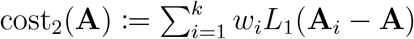.

*Proof*. Note that for any binary **A**, we have

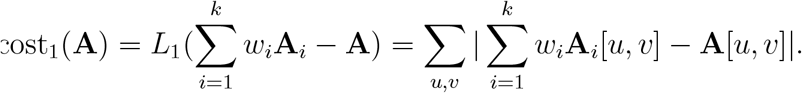

Since **A**_*i*_[*u, v*] ∈ [0, 1] and *w*_*i*_ ∈ [0, 1] and 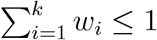, we have

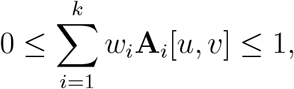

thus, we see that

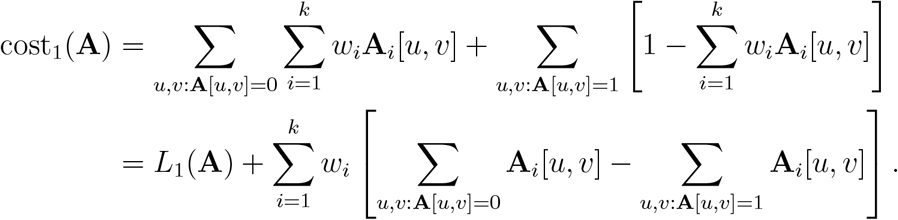

For cost_2_(**A**) we have

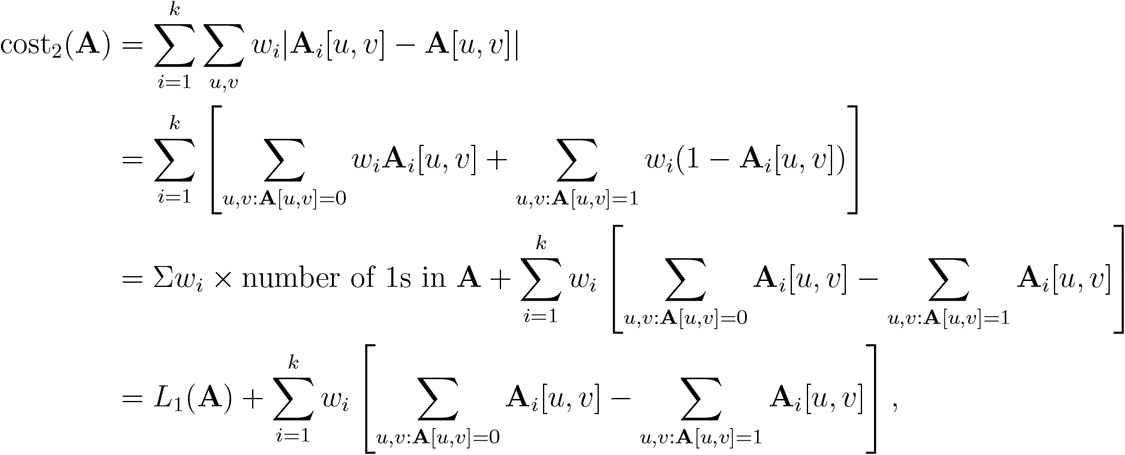

therefore, cost_2_(**A**) = cost_1_(**A**). □

##### Lemma 2.

Given a set of real matrices S = {**A**_1_, …, **A**_*k*_}, a weight vector **w** = {*w*_1_, …, *w*_*k*_} whose entries are also in [0, 1] and for which 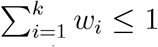, the following cost functions have identical minimizers over any subset of real matrices:

1. 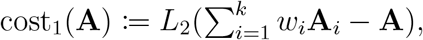,
2. 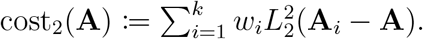.

*Proof*. Since cost_1_(**A**) is always non-negative, one may consider optimizing 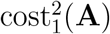. Expanding cost_1_(**A**) we have

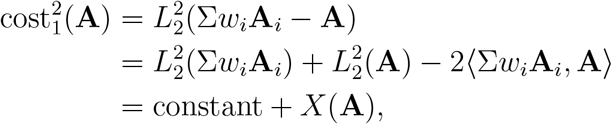

where,

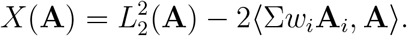

For cost_2_(**A**) we have

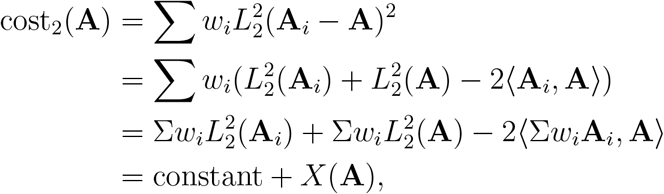

therefore, minimizing both cost_1_(**A**) and cost_2_(**A**) over any set reduces to minimizing *X*(**A**). □

#### Theorem 5.

Given any set of mutation trees *S* ⊂ *𝒯*_*n*_,

a. for any binary embedding *ε* : *𝒯*_*n*_ → {0, 1}^*n*×*n*^, the solution sets of 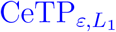 and 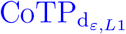 are identical.
b. for any embedding *ε*: *𝒯*_*n*_ → ℝ^*n*×*n*^, the solution sets of 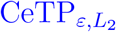 and 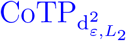 are identical.

*Proof*. Given a set of trees S = {*T*_1_, …, *T*_*k*_}, by setting **A**_**i**_ = *ε*(*T*_*i*_) and A = {**adj**(*T*) | *T* ∈ *𝒯*_*n*_},

a. the proof is straightforward using lemma 1 by minimizing costs over the set A.
b. the proof is straightforward using lemma 2 by minimizing costs over the set A.

### 5.4 Nearest Mapped Tree

#### 5.4.1 NMTP_adj,*L*_

##### Algorithm 2: TrimArb

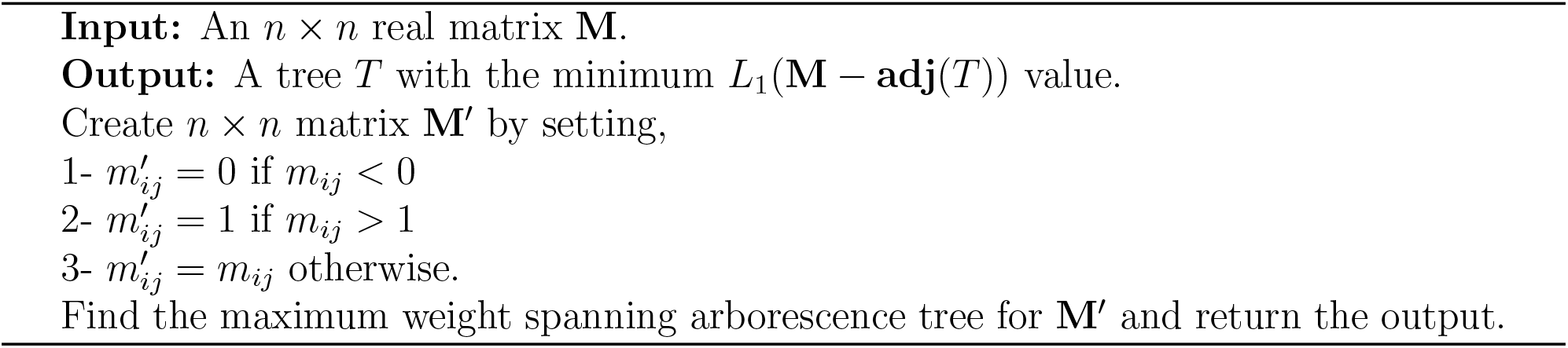

##### Theorem 6.

Given a real *n* × *n* matrix *M*,

a. the problem 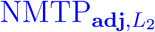 can be solved in *𝒪*(*n*) time through finding a maximum weight spanning arborescence tree.
b. the problem 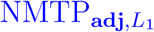 can be solved in O *𝒪*(*n*^3^) time by TrimArb algorithm (Supplementary Algorithm 2).

*Proof*.

a. The *Chu–Liu/Edmonds algorithm* [CL65; Edm67] can be used to find a spanning arborescence of minimum (or maximum) weight. Its running time is of *𝒪*(*V E*) and since all weights may be nonzero, its worst case running time is *𝒪*(*n*^3^). We treat **M** as an adjacency matrix of a weighted graph. Its maximum weight spanning arborescence *T*_max_ can be computed using Edmond’s algorithm. Set **A**^*^ = **adj**(*T*_max_). We claim that **A**^*^ is the optimal matrix. Assume *T* is a rooted tree on ℤ_*n*_ with **A**_*T*_ = **adj**(*T*). For the *L*_2_ norm we have

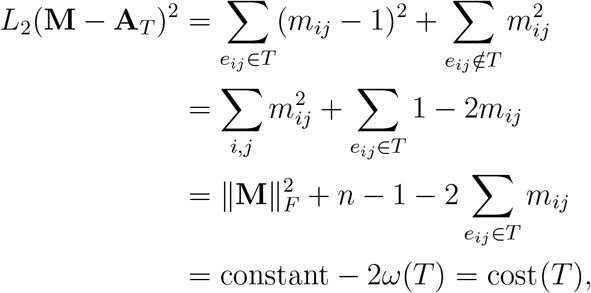

where 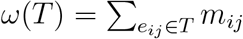 is the weight of *T*. The tree *T*_max_ has the largest *ω*(*T*) among trees on ℤ_*n*_, hence it is a minimizer of cost(*T*).
b. We treat **M** as an adjacency matrix of a weighted graph. Define three sets of edge indices as follows:

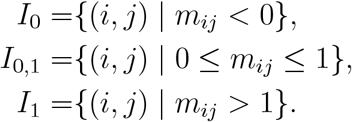

Assume that *T* is a rooted tree on ℤ_*n*_ with **A**_*T*_ = **adj**(*T*). For the *L*_1_ norm we have

*L*_1_(**M** − **A**_*T*_) =

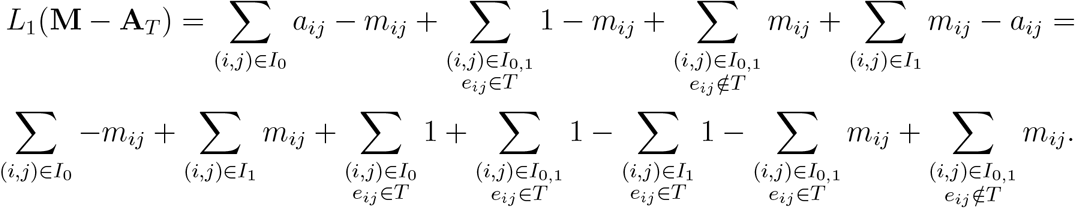

By adding and subtracting 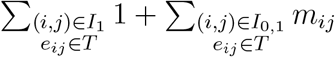 we have

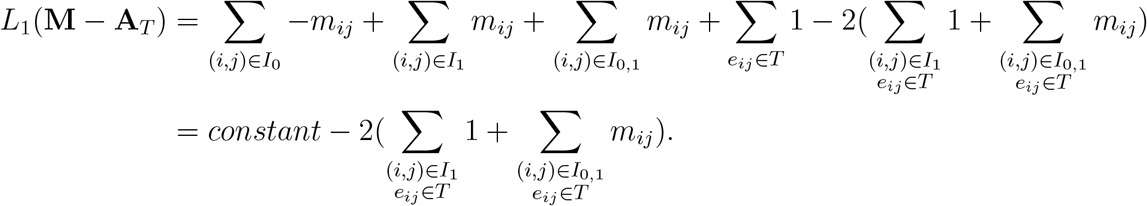

This means that in order to minimize *L*_1_(**M** − **A**_*T*_), the weight of tree edges in *I*_0,1_ is important, but for *I*_0_ and *I*_1_, just the number of tree edges in them matters. Tree edges in *I*_0_ enforce no added cost but tree edges in *I*_1_ reduces maximum possible cost for an edge which is 1. Therefore, the Algorithm TrimArb finds the optimal tree.

#### 5.4.2 NMTP_anc,*L*_

Firstly, we provide a set of necessary and sufficient conditions for a matrix to be an ancestry matrix of a rooted tree.

##### Lemma 3.

Matrix **X** is an ancestry matrix of a rooted tree *if and only if* all of the following constraints are satisfied:

1. The diagonal elements are zero.
2. There is a row with all entries equal to one, except the diagonal element.
3. For any two pairs of rows **x**_*i*_ and **x**_*j*_ one of the following conditions holds:
  - **x**_*i*_.**x**_*j*_ = 0 and **X**[*i, j*] = **X**[*j, i*] = 0.
  - 𝟙 (x_*i*_) ⊊ 𝟙 (x_*j*_) and **X**[*j, i*] =1.
  - 𝟙 (x_*j*_) ⊊ 𝟙 (x_*i*_) and **X**[*j, i*] =1.

*Proof*. Given a matrix **X** satisfying lemma 3’s constraints, we construct a tree *T* with the ancestry matrix **X**. First, we know that there is a row with all 1s but the diagonal element. It’s the root *r* of tree *T*. We go through the following steps repeatedly:

1. Pick a row **x**_*i*_ with the largest number of ones (among vertices not yet picked for the tree *T*).
2. Find a vertex **x**_*j*_ in *T* having the least number of 1s for which 𝟙 (**x**_*i*_) ⊂ 𝟙 (**x**_*j*_) and **X**[*j, i*] = 1.
3. Place **x**_*i*_ as a new child for **x**_*j*_ in *T*.

We know that there exists such **x**_*j*_ since 𝟙 (*r*) is a superset of ones for other rows and is the first vertex in the tree. We also know that **x**_*j*_ is unique since if we consider **x**_*k*_ and **x**_*l*_ such that **x**_*i*_ *⊂* **x**_*k*_ and **x**_*i*_ *⊂* **x**_*l*_ then **x**_*k*_.**x**_*l*_ = 0. Hence, either 𝟙 (**x**_*k*_) ⊊ 𝟙 (**x**_*l*_) or 𝟙 (**x**)_*l*_ ⊊ 𝟙 (**x**)_*k*_, implying that one of them has more 1s. This way every row of **X** is placed as a vertex in *T*.

Now, we want to prove that **X** is the ancestry matrix of *T*. Firstly, note that by the transitivity of the ⊊ operator, if *x*_*j*_ is an ancestor of *x*_*i*_, then 𝟙 (**x**_*i*_) ⊊ 𝟙 (**x**_*j*_). If **x**_*k*_ is the parent of **x**_*i*_ in *T*, by construction we know that **X**[*k, i*] = 1. Since **x**_*k*_ is either **x**_*j*_ itself or a descendant of it, we have 𝟙 (**x**_*k*_) ⊂ 𝟙 (**x**_*j*_), therefore **X**[*j, i*] = 1. It remains to show that if **X**[*j, i*] = 1, then **x**_*j*_ is an ancestor of **x**_*i*_ in *T*. We use induction on the depth of **x**_*i*_ in *T*. If the depth of **x**_*j*_ is 0, then it is the root vertex and there exist no *j* for which the condition is true, therefore the statement holds for the base case. Next, consider the case where the depth of **x**_*i*_ is *m* + 1. Since **X**[*j, i*] = 1 we know that 𝟙 (**x**_*i*_) ⊊ 𝟙 (**x**_*j*_). On the other hand, if **x**_*k*_ is the parent of **x**_*j*_, we have **x**_*j*_.**x**_*k*_ = 0, since **X**[*j, i*] = **X**[*k, i*] = 1. But **x**_*k*_ is the row with the least number of 1s which is a superset of **x**_*i*_, implying that 𝟙 (**x**_*k*_) ⊊ 𝟙 (**x**_*j*_) and **X**[*j, k*] = 1. Now, **x**_*k*_ is of depth *m* and by induction hypothesis we know that **x**_*j*_ is an ancestor of **x**_*k*_, hence the statement holds. This proves the *if* part. The *only if* part is rudimentary.

##### Theorem 7.

Given a matrix **M** ∈ ℝ^*n*×*n*^, the problem 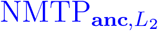 can be solved by AncL2 (Fig. 7).

*Proof*. In all of the following lemmas the matrix **X** is the optimal solution for AncL2.

**Note 5.1**. The cost function of the AncL2 ILP is

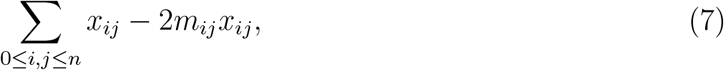

but the raw cost obtained by direct application of *L*_2_ norm is

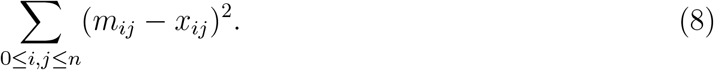

Since *m*_*ij*_s are constant and *x*_*ij*_s are 0 or 1, one may verify that

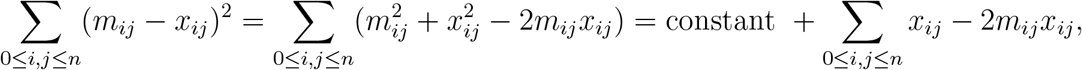

therefore minimizing 8 and 7 are the same problem.

**Note 5.2**. Any tree in *𝒯*_*n*_ would satisfy all of the constraints of AncL2 ILP, therefore the feasible region is not empty.

##### Lemma 4.

The graph *G* having the adjacency matrix **X** is transitive.

*Proof*. If there is an edge connecting *u* to *v* and an edge connecting *v* to *w*, then we know that *x*_*uv*_ = 1 and *x*_*vw*_ = 1. Therefore, the “transitivity” condition *x*_*uv*_ + *x*_*vw*_ − *x*_*uw*_ ≤ 1 implies that *x*_*uw*_ = 1 and there exist an edge between *u* and *w*. □

##### Lemma 5.

The graph *G* having the adjacency matrix **X** has no cycle.

*Proof*. First note that we have no self loop in *G* because for each vertex *i*, using the “no cycle” condition *x*_*ii*_ + *x*_*ii*_ ≤ 1, we have *x*_*ii*_ = 0. By Lemma 4 we know that *G* is transitive, therefore if we have a cycle of length greater than 1, for every pair of vertices *u* and *v* in the cycle we have *x*_*uv*_ = 1 and *x*_*vu*_ = 1 which contradicts the “no cycle” condition.

##### Lemma 6.

In the graph *G* having adjacency matrix **X**, if vertices *u* and *v* are connected to the vertex *k*, i.e., *x*_*uk*_ = *x*_*vk*_ = 1, then either *x*_*uv*_ = 1 or *x*_*vu*_ = 1 holds, but not both.

*Proof*. This is a direct consequence of the “common descendant” condition and lemma 5.

##### Lemma 7.

The matrix **X** which is a solution for AncL2 is an ancestry matrix.

*Proof*. For the proof we use Lemma 3.

1. Using Lemma 5 we know that all diagonal elements are zero.
2. Using the previous item and the “there should be a root” and the “just one root” conditions we know that there is a row in which all non-diagonal elements are equal to 1.
3. Considering two rows *u* and *v*,
4. if *x*_*uv*_ = *x*_*vu*_ = 0, then there is no *w* such that *x*_*uw*_ = *x*_*vw*_ = 1, since if there is such *w*, by Lemma 6 *x*_*uv*_ = 1 or *x*_*vu*_ = 1 which is a contradiction. Hence we have

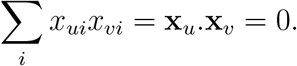
5. if *x*_*uv*_ = 1 and *x*_*vu*_ = 0 then by transitivity for each *w* where *x*_*vw*_ = 1, we have *x*_*uw*_ = 1, then

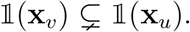
6. if *x*_*uv*_ = 0 and *x*_*vu*_ = 1 by a symmetrical argument we have

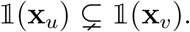
7. the case where *x*_*uv*_ = 1 and *x*_*vu*_ = 1 is not possible by Lemma 5. □

By Note 5.1 and Lemma 7 we see that the optimal solution to the AncL2 is the answer of the 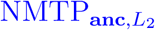 problem. □

**Note 5.3**. By introducing a condition of the form *x*_*ii*_ = 0 for all *i*’s, the two conditions “no cycle” and “just one root” may be omitted considering the “transivity” condition.

##### Theorem 8.

Given a matrix **M** ∈ R^*n*×*n*^, the problem 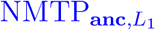 can be solved by AncL1 (Fig. 8).

*Proof*. The proof is almost the same as the proof for the Theorem 7. The cost function is different here and the variable *u*_*ij*_ is introduced to represent the value of |*x*_*ij*_ − *m*_*ij*_|. □

#### 5.4.3 NMTP_dist,*L*_

##### *𝒩𝒫*-hardness

###### Definition 6.

Partition Into Triangles (**PIT**): *Given a graph G* = (*V, E*) *where* |*V* | = 3*k for some k* ∈ ℕ *and each v* ∈ *V is covered by some triangles in G, can V be partitioned into disjoint sets V*_1_, …, *V*_*k*_ *each containing exactly 3 vertices, such that each of these V*_*i*_*s is the vertex set of a triangle in G?*

###### Theorem 9.

([GJ79]) The problem Partition Into Triangles is *𝒩𝒫*-complete.

**Note 5.4**. The original problem stated in [GJ79] does not have the condition that each vertex should be covered by some triangle in the given graph *G*. But adding this condition does not change the complexity since it is possible to be checked in polynomial time.

###### Theorem 10.

The problems 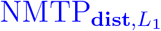 and 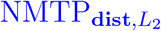 are *𝒩𝒫*-hard.

*Proof*. Given a graph *G* = (*V, E*) with |*V* | = *n* = 3*z* with *p* triangles, we want to reduce PIT to 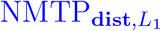 and 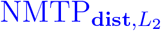. We have to create a matrix **M** based on *G* and solve 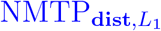 for **M** and create a triangulation of *V* if there exist one using this solution.

In order to create **M**, consider a graph *G*^′^ with vertices and edges in 5 layers described as,

1. Layer-R: A single vertex *r*.
2. Layer-S: A set of *n*^5^ vertices. Each vertex in this layer is denoted by *s*_*i*_. All of them are connected directly to *r*.
3. Layer-S2: The union of *n*^5^ sets of *n*^5^ vertices: For each *s*_*i*_ in Layer-S there are *n*^5^ vertices denoted by *s*_*ij*_ directly connected to it.
4. Layer-TR: A set of *p* vertices for each triangle tr_*i*_ in *G*. Each vertex denoted by *t*_*i*_, is directly connected to *r*.
5. Layer-V: The set *V*. Each vertex *v* in this layer is connected to every *t*_*i*_ for which *v* ∈ tr_*i*_.

**Observation 5.2**. The distances between different layers of vertices in *G*^′^ are described in the following table.

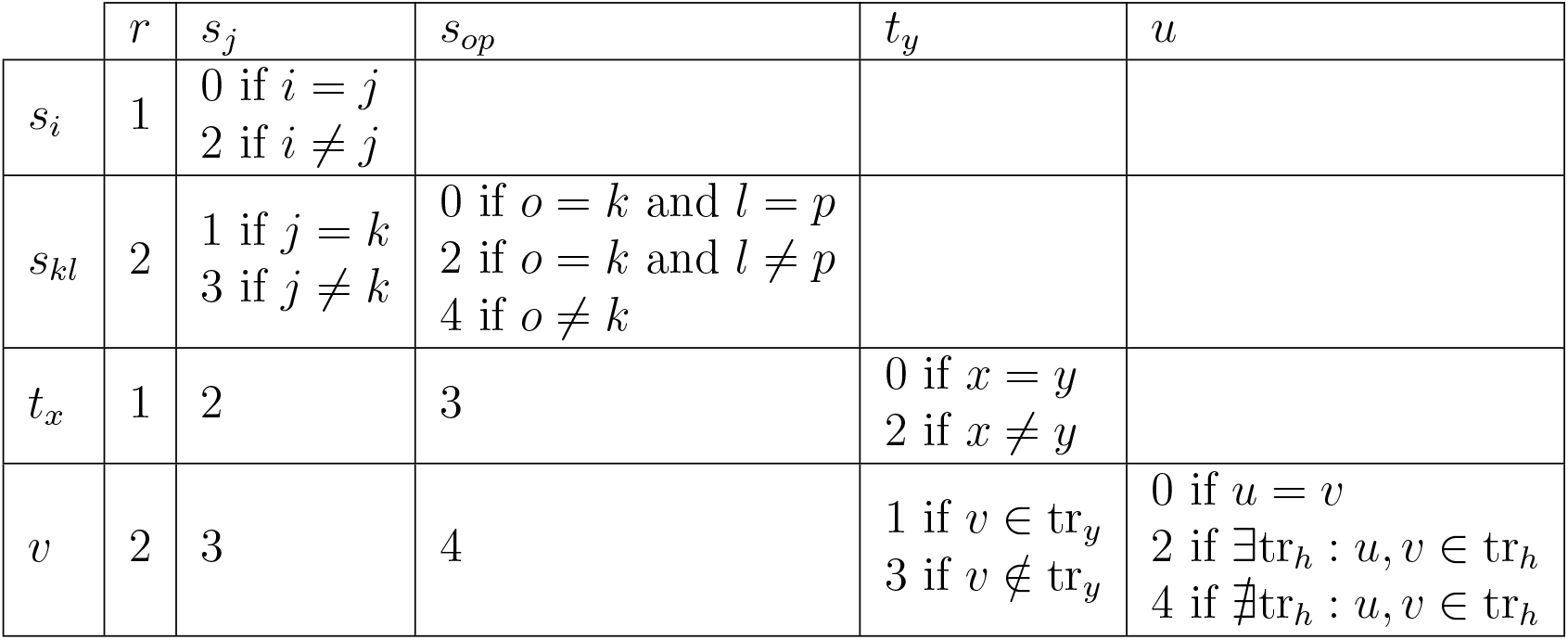

Now define the matrix **M** whose entries are identical to those of **dist**(*G*^′^) except for all (pairs of) vertices of Layer-V for which we define the entries as

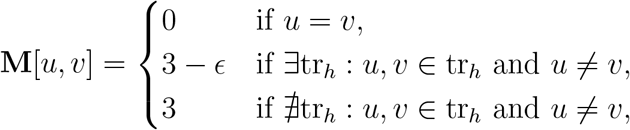

where *ϵ* is a small nonzero real number to be set later. Let *T* with distance matrix **D** be an optimal solution of 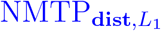 for **M**. It is obvious that *V* (*T*) = *V* (*G*^′^). We first want to show that *E*(*G*^′^) − *E*(*T*) ⊂ Layer-V × Layer-TR.

**Observation 5.3**. *We see that L*_1_(**M** − **D**) = *𝒪* (*n*^4^). Consider a tree *T*^′^ with all edges between vertices in layers Layer-R, Layer-S, Layer-S2, and Layer-TR from *G*^′^. Then we connect any *v* in Layer-V to exactly one *t*_*i*_ in Layer-TR. Set **D**^′^ = **dist**(*T* ^′^). In *T*^′^ the distance between vertices in three layers Layer-R, Layer-S, and Layer-S2 to all vertices in *G*^′^ are the same as in **M**. Furthermore the distance between vertices in Layer-TR with each other, is also consistent with **M**. The only distances that may not match with **M** are distances between vertices in Layer-V and vertices in Layer-V or Layer-TR. Because there are *p* triangles in *G*, we have 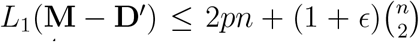. Since *p* = *𝒪* (*n*^3^), we have *L*_1_(**M** − **D**) ≤ *L*_1_(**M** − **D**^′^) = *𝒪* (*n*^4^).

**Observation 5.4**. *In the tree T, the vertex r remains directly connected to all vertices in Layer-S*. By observation 5.3, *r* remains connected to O(*n*^5^) vertices from Layer-S. Now suppose that there exist a vertex *s*_*i*_ from Layer-S, for which (*r, s*_*i*_) ∉ *E*(*T*). Since *T* is a tree, there is just one path between *r* and *s*_*i*_. If we remove one edge in this path, *s*_*i*_ becomes disconnected from *r*. Since we just removed one vertex, *r* is still connected directly to *𝒪* (*n*^5^) vertices from Layer-S. This means the distance between *s*_*i*_ and *𝒪* (*n*^5^) vertices from Layer-S is at least 3, which makes *L*_1_(**M** − **D**) = *Ω* (*n*^5^) which is a contradiction.

**Observation 5.5**. *In the tree T, all vertices in Layer-S2 remain directly connected to the corresponding vertices from Layer-S*. The proof is similar to observation 5.4.

**Observation 5.6**. *In the tree T, all vertices in Layer-TR remain directly connected to r*. Vertex *r* is directly connected to all vertices in Layer-S. A similar argument to observation 5.4 reveals that the distance for all vertices in Layer-TR to *r* should remain 1.

**Observation 5.7**. *In the tree T, all vertices in Layer-V remain directly connected to vertices in Layer-TR*. Consider a vertex *v* from Layer-V. If it is connected directly to a vertex from Layer-S2 like *s*_*ij*_, then its distance to Ω(*n*^5^) vertices connected to *s*_*i*_ reduces from 4 to 3. If it is directly connected to a vertex from Layer-S like *s*_*i*_, then its distance to Ω(*n*^5^) vertices connected to *s*_*i*_ reduces from 4 to 2. If it is directly connected to the single vertex in Layer-R namely *r*, then its distance to Ω(*n*^5^) vertices connected to *r* reduces from 3 to 2. Thus *v* cannot be connected directly to any vertex in any of the Layer-R, Layer-S or Layer-S2. It also cannot be connected directly to any other vertex *u* from Layer-V, because it makes their distance unequal to all other vertices in the tree, because for any vertex in *T*, it is either closer to *v* or *u*. Considering just vertices in Layer-S2, if *v* is directly connected to *u*, then *L*_1_(**M** − **D**) = Ω(*n*^10^). Therefore, vertices in Layer-V are directly connected to vertices in Layer-TR.

**Observation 5.8**. *In the tree T, each vertex v from Layer-V is directly connected to one vertex in Layer-TR*. Since all vertices in Layer-TR are directly connected to *r, v* cannot be connected to two vertices in Layer-TR.

Now we want to show that if there exist a triangulation of *G*, the vertices from Layer-V, are arranged in *T* in a way that this triangulation can be easily constructed.

**Observation 5.9**. *In the tree T, each v from Layer-V is connected to a t*_*i*_ *from Layer-TR for which v* ∈ *tr*_*i*_. Suppose *v* ∉ tr_*i*_, then let *T*^′^ be a tree obtained by disconnecting *v* from *t*_*i*_ and connect it to some *t*_*j*_ for which *v* ∈ tr_*j*_. Defince *X*(*x*) = *u* ∈ *V* | *u* is connected to *t*_*x*_}. Then the distances of *T* and *T* ^′^ are identical except for distances between *v* and vertices in ({*t*_*i*_, *t*_*j*_} ∪ *X*(*i*) ∪ *X*(*j*)) − {*v*}. The difference of sum of the distances between *v* and

*X*(*i*) ∪ *X*(*j*) in *T* and *T* ^′^ for *L* ∈ {*L*_1_, *L*_2_} is

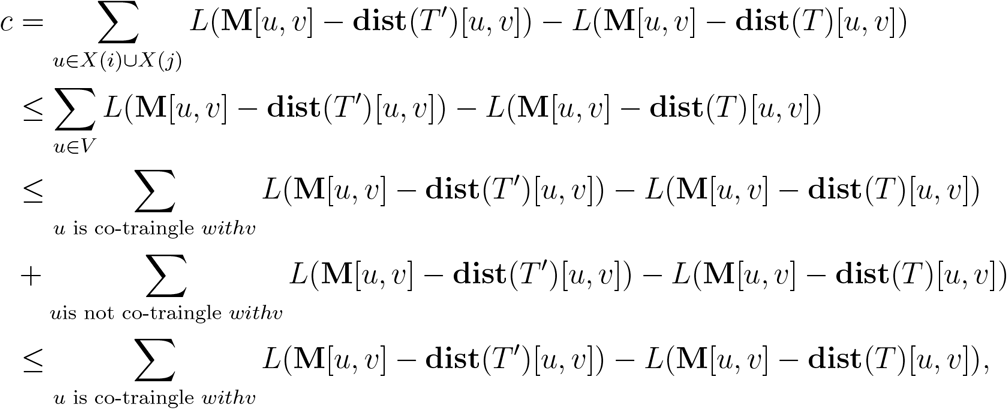

since if *u* is not co-triangle with *v* for

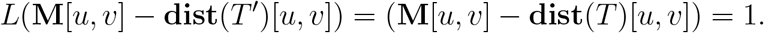

Now, for *L*1 we have

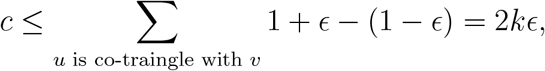

and for *L*2 we have

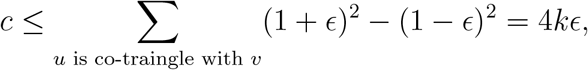

where *k* is the number of vertices which are co-triangle with *v*. As a result, if we choose *ϵ* small enough, *c* is bounded by 1. Hence, the distance between *v* and *t*_*i*_ is increased from 1 to 3 and distance between *v* and *t*_*j*_ is decreased from 3 to 1. This means that **dist**(*T* ^′^) is closer to the **M** from **D**, which is a contradiction.

**Observation 5.10**. *If every vertex v from Layer-V in T has exactly two other siblings, then this gives a triangulation of G*. By Observations 5.8 and 5.9, if one considers the sets *V*_*i*_ = {*v*|*v* ∈ Layer-V and (*v, t*_*i*_) ∈ *E*(*T*)}, there are 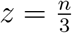 nonempty *V*_*i*_ which are all disjoint and each of them having exactly 3 members and covering corresponding triangle *tr*_*i*_.

**Observation 5.11**. *If there is a triangulation of V, then T will specify one*. If for every *v* in Layer-V, it has two siblings from Layer-V connected to the same *t*_*i*_, then, *T* gives us a triangulation. If *T* is not giving a triangulation, suppose *T* ^′^ is a tree obtained by one of the triangulations of *G*. By observation 5.9 we know that in *T* every vertex *v* from Layer-V has at most two siblings. The distances between all vertices in all layers except Layer-V are the same for both *T* and *T* ^′^. The distances between Layer-V and three layers Layer-R, Layer-S and Layer-S2 is the same too for both trees. The distances between Layer-V and Layer-TR may be different for some vertices. But the difference between distances sums up to zero, because every vertex is connected to exactly one of it’s covering triangles. It has distance 1 to it and distance 3 to all other triangles.

Now, consider the difference between distances of vertices in Layer-V in the input matrix **M**. For a vertex *v* in *T* with *q* siblings (0 ≤ *q* ≤ 2), in the case of *L*_1_ norm we have

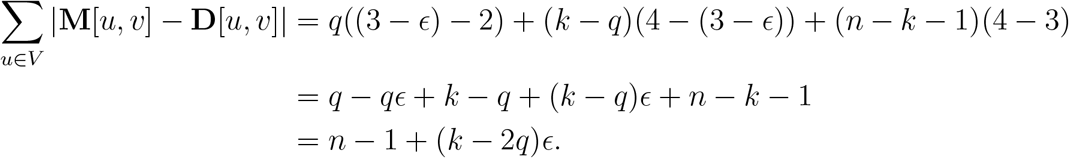

For *T* ^′^ we have

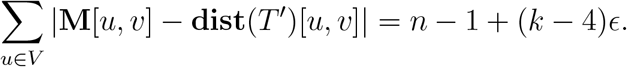

For a vertex *v* in *T* with *q* siblings (0 ≤ *q* ≤ 2), in the case of *L*_2_ norm we have

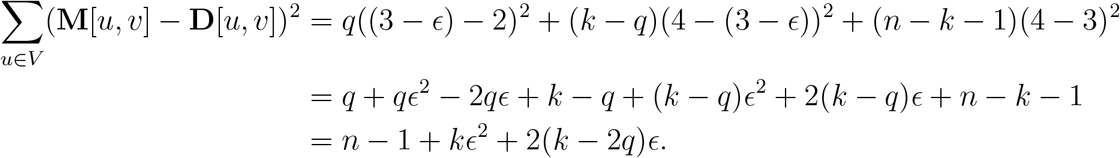

For *T* ^′^ and norm *L*_1_ we have

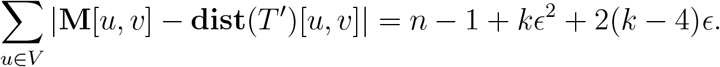

The tree *T* does not impose a triangulation, therefore there exist a *v* in Layer-V with *q <* 2, accordingly for *L* ∈{*L*_1_, *L*_2_} we have *L*(**M** − **dist**(*T*^′^)) *< L*(**M** − **D**) which is a contradiction.

We used the parameter *ϵ* in Observations 5.9 and 5.11 and it is straight forward to verify that If one chooses

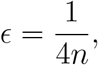

then both observations are true.

##### ILPs

###### Theorem 11.

Given a matrix **M** ∈ ℝ^*n*×*n*^, the ILP program DistL1 solves 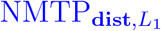.

*Proof*. The solution to the DistL1 is a graph *G* with the adjacency matrix *E* for which *E*[*i, j*] = *e*_*ij*_. First, note that *E* is symmetric, irreflexive and has *n* − 1 undirected edges. Thus, in order to prove that it is a tree with minimum distance from **M**, it suffices to show that it has no cycles and also **D** is its distance matrix.

**Note 5.5**. Any tree in *𝒯*_*n*_ would satisfy all of the constraints of AncL2 ILP, therefore the feasible region is not empty.

**Observation 5.12**. *Variables e*_*ij*_ = 1 *iff d*_*ij*_ = 1. If *d*_*ij*_ = 1 then by condition *e*_*ij*_ + *d*_*ij*_ ≥ 2 we have that *e*_*ij*_ = 1. If *d*_*ij*_ *>* 1 then Σ_*k*_ *z*_*ijk*_ *>* 0 and thus *K* = {*k*|*z*_*ijk*_ = 1} is not empty. Suppose *z*_*ijk*_ = 1 then by condition *z*_*ijk*_ + *z*_*ikj*_ + *z*_*jki*_ ≤ 1 and as a result *z*_*ikj*_ = *z*_*jki*_ = 0. Now by condition *e*_*ij*_ − *z*_*ikj*_ − *z*_*jki*_ ≤ 0 we know that *e*_*ij*_ = 0.

**Observation 5.13**. *Graph G is connected and every pair of vertices u and v are connected by a path of length d*_*uv*_. We know that *d*_*uv*_ ≥ 1. Let us use strong induction for the proof. For the base case, if *d*_*uv*_ = 1 then by observation 5.12 we know that there is an edge between them, therefore they are connected with a path of length 1. Now assume that for *d* = 1 to *m* the statement holds. If *d*_*uv*_ = *m* + 1 then by condition “vertices between *i* and *j*” there exist *k* for which *z*_*uvk*_ = 1. By “triangle equality” condition we know *d*_*uv*_ ≤ *d*_*uk*_ + *d*_*kv*_ and by “additive distance” condition we know *d*_*uv*_ ≥ *d*_*uk*_ + *d*_*kv*_, therefore *d*_*uv*_ = *d*_*uk*_ + *d*_*kv*_. Since *d*_*uk*_ ≤ *m* and *d*_*kv*_ ≤ *m*, by induction hypothesis there exist a path of length *d*_*uk*_ from *u* to *k* and a path of length *d*_*kv*_ from *k* to *v*. Hence, *u* and *v* are connected by a path of length *d*_*uv*_ = *d*_*uk*_ + *d*_*kv*_ passing through *k*.

This proves that *G* is indeed a tree, since it has *n* − 1 edges and is connected. Also we proved that **dist**(*G*)[*u, v*] = *d*_*uv*_. Any tree satisfies all of these conditions, therefore the tree obtained by solving this ILP is a solution to 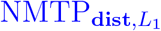.

###### Theorem 12.

Given a matrix **M** ∈ ℝ^*n*×*n*^, the problem 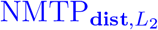 can be solved by DistL2 (Fig. 11).

###### Theorem 13.

Given a matrix **M** ∈ ℝ^*n*×*n*^, the problem 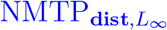 can be solved by DistLInf (*Fig*.10).

*Proof*. The variable *u* here is an upper bound for *L*_1_(*M* − *X*). Other parts of the proof is the same as Theorem **??**.

### 5.5 Weighted Centroids

#### Definition 7

(The Weighted Consensus Tree Problem (WCoTP_*d*,*w*_)). ([GSO23]) Let *d* : *𝒯*_*n*_ × *𝒯*_*n*_ → ℝ be a dissimilarity measure, set 𝒮 = {*T*_1_, …, *T*_*k*_} ⊂ T_*n*_ be a set of mutation trees, and *w* : 𝒮 → [0, 1] a weighting function with 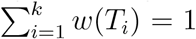. Then the *weighted consensus tree* of 𝒮, denoted by *𝒯*_con_ ∈ *𝒯*_*n*_, is defined as

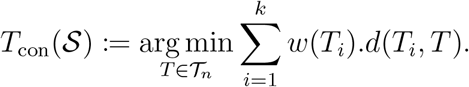

**Note 5.6**. Removing the condition 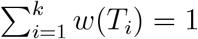 on weight does not expand the solutions to the weighted consensus tree problem. If we consider weights *w* with 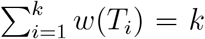, it has the same solution as 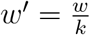 for which we have 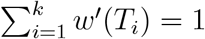.

#### Theorem 14.

The weighted centroid tree problem 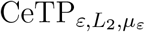 is a generalization of weighted consensus tree problem 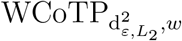, in the sense that for every weighting function *w* for 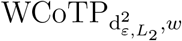, the 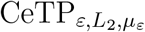 defined for the weighted sum operator *μ*_*ε*_ with weight vector *w*, has the same solution set.

*Proof*. Given a set of trees S = {*T*_1_, …, *T*_*k*_} ⊂ T_*n*_ and the 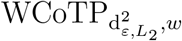 problem, consider 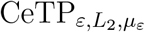 with *w* as the weights for *μ*_*ε*_. Expanding the cost function of the weighted centroid tree problem for a tree *T* we have

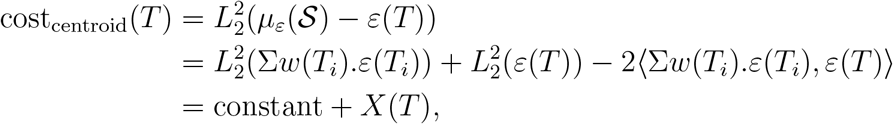

where

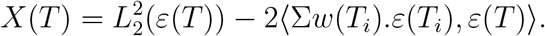

For the consensus tree we have

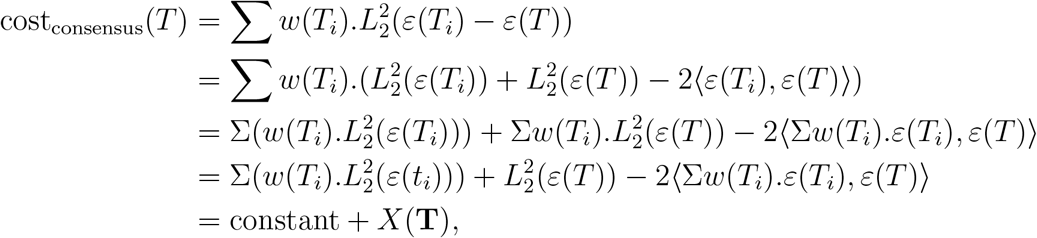

therefore, minimizing both of the cost_centroid_(*T*) and cost_consensus_(*T*) reduces to minimizing *X*(*T*).

**Note 5.7**. It ought to be noted that the problem 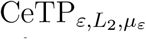 is an absolute generalization of the problem 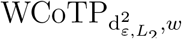, meaning that also for every weight *w* for 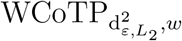, there exist a weighted sum operator *μ*_*ε*_ for which 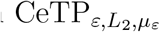 has the same solution set, the inverse is not true. The most straightforward example is 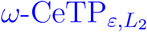, for which the weights are uniform but 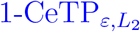, 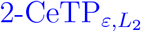, and 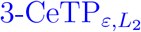 has different solution spaces. As noted in Note 5.6, scaling the weights in 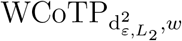 does not change the solution space.

### 5.6 Simulation Method

The function `random-tree` from the NetworkX(2.6.3) package, with default parameters, is used to generate random trees. For reproducibility, a seed parameter is used for every random function involved in generating and altering trees. The generation of CCfs (Cancer Cell Fraction) is done in a top-down manner. The CCF for root vertex is 1. To generate the CCFs for *k* children of a vertex *v* with known CCF, we generate *k* + 1 uniform random numbers in [0, 1] and normalize them so that their sum is equal to CCF(*v*) and assign first *k* normalized values to the children. Three types of possible alterations can be applied to the generated ground-truth trees:

1. **pc** (parent-child): This involves swapping a child vertex with its parent. Th substitu-tion is perforemed if

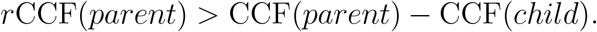 Where *r* is a random number in [0, 1]. The child vertex is uniformly picked from the vertices of the tree. We also swap the CCFs for the parent and child vertices so that CCF values remain consistent with the new structure of the tree.
2. **bm** (branch-move): This involves moving a vertex with all of its descendants to a vertex which has enough *empty CCF*. The empty CCF of vertex *v* is defined as

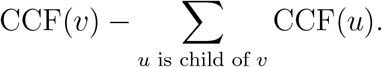 The vertex *v* is uniformly selected from tree vertices. Then possible parents, i.e., vertices with enough empty CCF are computed. If the set of possible parents is not empty, one of them is selected uniformly as the new parent.
3. **nr** (node-remove): This involves removing a vertex from its place in the tree and placing it as a direct child of the root vertex. Its former parent becomes the new parent of its former children. Firstly, vertex *v* is selected uniformly from the tree vertices. Then it will become a direct child of the root vertex if

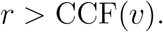

where *r* is a random number in [0, 1]. After moving the vertex *v*, we set CCF(*v*) = *ϵ* because the root vertex may not have any empty CCF. We used *ϵ* = 10^−6^. A parameter **cp** (change-probability) is used to determine the number of alterations applied to each tree. For each alteration, we repeatedly generate a random number *r* ∈ [0, 1]. If *r >* **cp** we apply the alteration. If not, we stop applying this type of alteration and move on to the next type of alteration. The order of application of alterations is the same as listed above. The source code is also available at the git repository associated with this paper.

### 5.7 Supplementary images and tables

In all images, the algorithm AncstILP2 is the same as WAncILP_2_ and AncstILP3 is the same as WAncILP_3_.

## References

[AL20] Nancy G. Azizian and Yulin Li. “XPO1-dependent nuclear export as a target for cancer therapy”. In: Journal of Hematology & Oncology 13.1 (Dec. 2020), p. 61.

[AQE19] Nuraini Aguse, Yuanyuan Qi, and Mohammed El-Kebir. “Summarizing the solution space in tumor phylogeny inference by multiple consensus trees”. In: Bioinformatics. Vol. 35. 14. July 2019, pp. i408–i416.

[Art88] Spyros Artavanis-Tsakonas. “The molecular biology of the Notch locus and the fine tuning of differentiation in Drosophila”. In: Trends in Genetics 4.4 (Apr. 1988), pp. 95–100.

[BBG20] Giulia Bernardini, Paola Bonizzoni, and PaweÅ Gawrychowski. “On Two Measures of Distance between Fully-Labelled Trees”. In: Leibniz International Proceedings in Informatics, LIPIcs 161 (2020), pp. 1–17. arXiv: 2002.05600.

[Ber+19] Giulia Bernardini et al. “A rearrangement distance for fully-labelled trees”. In: Leibniz International Proceedings in Informatics, LIPIcs 128.23 (2019). arXiv: 1904.01321.

[Bry03] David Bryant. “A classification of consensus methods for phylogenetics”. In: 0000 (2003), pp. 163–183.

[Car+18] Giulio Caravagna et al. “Detecting repeated cancer evolution from multi-region tumor sequencing data”. In: Nature Methods 15.9 (2018), pp. 707–714.

[Chr+20] Sarah Christensen et al. “Detecting evolutionary patterns of cancers using consensus trees”. In: Bioinformatics 36 (2020), pp. I684–I691.

[Cic+21] Simone Ciccolella et al. “Triplet-based similarity score for fully multilabeled trees with poly-occurring labels”. In: Bioinformatics 37.2 (2021), pp. 178–184.

[CL65] Yeong-Jin; Chu and Tseng-Hong Liu. “On the shortest arborescence of a directed graph”. In: Scientia Sinica 14 (1965), pp. 1396–1400.

[Cra+22] Karen Cravero et al. “NOTCH1 PEST domain variants are responsive to standard of care treatments despite distinct transformative properties in a breast cancer model”. In: Oncotarget 13 (2022), pp. 373–386.

[DiN+20] Zach DiNardo et al. “Distance measures for tumor evolutionary trees”. In: Bioinformatics 36.7 (Apr. 2020). Ed. by Yann Ponty, pp. 2090–2097.

[Edm67] Jack Edmonds. “Optimum branchings”. In: Journal of Research of the National Bureau of Standards Section B Mathematics and Mathematical Physics 71B.4 (Oct. 1967), p. 233.

[El-+16] Mohammed El-Kebir et al. “Inferring the Mutational History of a Tumor Using Multi-state Perfect Phylogeny Mixtures”. In: Cell Systems 3.1 (2016), pp. 43–53.

[FS21] Xuecong Fu and Russell Schwartz. “ConTreeDP: A consensus method of tumor trees based on maximum directed partition support problem”. In: Proceedings - 2021 IEEE International Conference on Bioinformatics and Biomedicine, BIBM 2021 Mcmc (2021), pp. 125–130.

[FV90] Eric R. Fearon and Bert Vogelstein. “A genetic model for colorectal tumorigenesis”. In: Cell 61.5 (June 1990), pp. 759–767.

[GJ79] Micheal R. ; Garey and David S. Johnson. Computers and Intractability: A Guide to the Theory of NP-Completeness (Series of Books in the Mathematical Sciences). First Edit. W.H. Freeman, 1979.

[Gov+20] Kiya Govek et al. “GraPhyC: Using Consensus to Infer Tumor Evolution”. In: IEEE/ACM Transactions on Computational Biology and Bioinformatics 19.1 (Jan. 2020), pp. 465–478.

[GSO18] Kiya Govek, Camden Sikes, and Layla Oesper. “A Consensus Approach to Infer Tumor Evolutionary Histories”. In: Proceedings of the 2018 ACM International Conference on Bioinformatics, Computational Biology, and Health Informatics. New York, NY, USA: ACM, Aug. 2018, pp. 63–72.

[GSO23] Ziyun Guang, Matthew Smith-Erb, and Layla Oesper. “A weighted distance-based approach for deriving consensus tumor evolutionary trees”. In: Bioinformatics 39. Supplement 1 (June 2023), pp. i204–i212.

[HBD10] Kasper D. Hansen, Steven E. Brenner, and Sandrine Dudoit. “Biases in Illumina transcriptome sequencing caused by random hexamer priming”. In: Nucleic Acids Research 38.12 (July 2010), e131–e131.

[HG18] Joanna Huszno and Ewa Grzybowska. “TP53 mutations and SNPs as prognostic and predictive factors in patients with breast cancer”. In: Oncology Letters 16.1 (2018), pp. 34–40.

[Hod+20] Ermin Hodzic et al. “Identification of conserved evolutionary trajectories in tumors”. In: Bioinformatics 36 (2020), pp. I427–I435.

[HW11] Douglas Hanahan and Robert A. Weinberg. “Hallmarks of cancer: The next generation”. In: Cell 144.5 (2011), pp. 646–674.

[JBZ21] Katharina Jahn, Niko Beerenwinkel, and Louxin Zhang. “The Bourque distances for mutation trees of cancers”. In: Algorithms for Molecular Biology 16.1 (Dec. 2021), p. 9.

[Jia+20] Tao Jiang et al. “ALOX12B promotes carcinogenesis in cervical cancer by regulating the PI3K/ERK1 signaling pathway”. In: Oncology Letters 20.2 (2020), pp. 1360–1368.

[Kar+19] Nikolai Karpov et al. “A multi-labeled tree dissimilarity measure for comparing “lonal trees” of tumor progression”. In: Algorithms for Molecular Biology 14.1 (2019), pp. 1–18.

[Kha+19] Sahand Khakabimamaghani et al. “Collaborative intra-tumor heterogeneity detection”. In: Bioinformatics. Vol. 35. 14. 2019, pp. i379–i388.

[KMF01] Peter Krieg, Friedrich Marks, and Gerhard Fürstenberger. “A Gene Cluster Encoding Human Epidermis-type Lipoxygenases at Chromosome 17p13.1: Cloning, Physical Mapping, and Expression”. In: Genomics 73.3 (May 2001), pp. 323–330.

[KMS22] Ekaterina Kim, Daria A. Mordovkina, and Alexey Sorokin. “Targeting XPO1-Dependent Nuclear Export in Cancer”. In: Biochemistry (Moscow) 87.S1 (Jan. 2022), S178–S191.

[Lan+23] Jennifer R. Landes et al. “The efficacy of selinexor (KPT-330), an XPO1 inhibitor, on non-hematologic cancers: a comprehensive review”. In: Journal of Cancer Research and Clinical Oncology 149.5 (May 2023), pp. 2139–2155.

[Lee+09] Ji Young Lee et al. “Candidate gene approach evaluates association between innate immunity genes and breast cancer risk in Korean women”. In: Carcino-genesis 30.9 (2009), pp. 1528–1531.

[LK06] Kevin G. Leong and Aly Karsan. “Recent insights into the role of Notch signaling in tumorigenesis”. In: Blood 107.6 (Mar. 2006), pp. 2223–2233.

[LKB23] Xiang Ge Luo, Jack Kuipers, and Niko Beerenwinkel. “Joint inference of exclusivity patterns and recurrent trajectories from tumor mutation trees”. In: Nature Communications 14.1 (2023), pp. 1–14.

[LOA11] Camille Lobry, Philmo Oh, and Iannis Aifantis. “Oncogenic and tumor suppressor functions of Notch in cancer: it’s NOTCH what you think.” In: The Journal of experimental medicine 208.10 (Sept. 2011), pp. 1931–5.

[LRV20] Mercè Llabrés, Francesc Rosselló, and Gabriel Valiente. “A Generalized Robinson-Foulds Distance for Clonal Trees, Mutation Trees, and Phylogenetic Trees and Networks”. In: Proceedings of the 11th ACM International Conference on Bioinformatics, Computational Biology and Health Informatics, BCB 2020 (2020).

[LRV21] Mercè Llabrés, Francesc Rossello, and Gabriel Valiente. “The Generalized Robinson-Foulds Distance for Phylogenetic Trees”. In: Journal of Computational Biology 28.12 (2021), pp. 1181–1195.

[Mat+17] Yusuke Matsui et al. “phyC: Clustering cancer evolutionary trees”. In: PLoS Computational Biology 13.5 (2017), pp. 1–17.

[MP10] Andriy Marusyk and Kornelia Polyak. “Tumor heterogeneity: Causes and consequences”. In: Biochimica et Biophysica Acta (BBA) - Reviews on Cancer 1805.1 (Jan. 2010), pp. 105–117.

[PAJ13] Trang T. Pham, Steven P. Angus, and Gary L. Johnson. “MAP3K1: Genomic Alterations in Cancer and Function in Promoting Cell Survival or Apoptosis”. In: Genes and Cancer 4.11-12 (2013), pp. 419–426.

[Par+16] Swati Parekh et al. “The impact of amplification on differential expression analyses by RNA-seq”. In: Scientific Reports 6.1 (May 2016), p. 25533.

[PV22] Leonardo Pellegrina and Fabio Vandin. “Discovering significant evolutionary trajectories in cancer phylogenies”. In: Bioinformatics 38.Supplement 2 (Sept. 2022), pp. ii49–ii55.

[QE23] Yuanyuan Qi and Mohammed El-Kebir. “Consensus Tree under the Ancestor-Descendant Distance is NP-hard”. In: bioRxiv (Jan. 2023), p. 2023.07.17.549375.

[Raz+18] Pedram Razavi et al. “The Genomic Landscape of Endocrine-Resistant Advanced Breast Cancers”. In: Cancer Cell 34.3 (2018), 427–438.e6.

[RF81] D.F. Robinson and L.R. Foulds. “Comparison of phylogenetic trees”. In: Mathematical Biosciences 53.1-2 (Feb. 1981), pp. 131–147.

[Roo+15] Michael S Rooney et al. “Molecular and genetic properties of tumors associated with local immune cytolytic activity.” In: Cell 160.1-2 (Jan. 2015), pp. 48–61.

[She+09] Min Shen et al. “Polymorphisms in innate immunity genes and lung cancer risk in Xuanwei, China”. In: Environmental and Molecular Mutagenesis 50.4 (May 2009), pp. 285–290. doi: 10.1002/em.20452.

[You+22] Hassan Yousefi et al. “Notch signaling pathway: a comprehensive prognostic and gene expression profile analysis in breast cancer”. In: BMC Cancer 22.1 (2022), pp. 1–21.

